# Overexpression and nonsynonymous mutations of UDP-glycosyltransferases potentially associated with pyrethroid resistance in *Anopheles funestus*

**DOI:** 10.1101/2023.08.25.554687

**Authors:** Talal Al-Yazeedi, Abdullahi Muhammad, Helen Irving, Seung-Joon Ahn, Jack Hearn, Charles S. Wondji

## Abstract

UDP-glycosyltransferases (UGTs) constitute a superfamily of enzymes that play a vital role in the biotransformation of diverse hydrophobic substrates into more hydrophilic products, thereby facilitating their excretion from the cell through transporters. The significance of UGTs in conferring insecticide resistance has been emphasized in various insect species. In this study, we characterised *Anopheles funestus* UGT genes genome-wide and explored their evolution and association with pyrethroid resistance. We combined genome-wide association of pooled-template sequencing (GWAS-PoolSeq) with the transcriptomic profile of pyrethroid-resistant *An. funestus* populations, and deep targeted sequencing of UGTs from 80 individual mosquitoes collected in Malawi, Uganda, Cameroon and the two laboratory colonies (FANG and FUMOZ) to investigate the role of UGTs in pyrethroid resistance. We identified common overexpression of UGT310B2 (AFUN000679) in the resistant laboratory colony (FUMOZ) and resistant field populations from Malawi, Cameroon and Uganda. Significant gene-wise *F_st_* differentiation between the resistant and putatively susceptible populations was observed for UGT301C2 and UGT302A3 in Malawi, as well as UGT306C2 in Uganda. Furthermore, the gene-wise Tajimas D density curves of the sequenced regions provided insights into genome-wide processes elucidating population structures within *An. funestus* populations from these three countries, supporting previous observations. Additionally, we identified significantly differentiated nonsynonymous mutations within UGT genes, which may potentially contribute to pyrethroid resistance. The identified role of *An. funestus* UGT genes in pyrethroid resistance has direct implications for current vector control strategies, management approaches, and the prediction of potential cross-resistance to other insecticides that can be directly detoxified by UGTs.

## 1. Introduction

In sub-Saharan Africa, malaria remains a significant cause of morbidity and mortality, particularly among pregnant women and children under 5 years old. It accounts for more than 96% of malaria-related deaths globally [1]. The primary strategies for malaria control depend heavily on insecticide-based interventions, such as indoor residual spray (IRS) and long-lasting insecticide nets (LLINs) [1, 2]. These interventions have been highly successful in reducing malaria cases and associated morbidity, globally preventing over 2 billion malaria cases and 11.7 million malaria-related deaths between 2000 and 2021 [1]. However, several mosquito species have developed multiple and cross-resistance to several insecticides, including pyrethroids the main compound approved by WHO for LLINs. Unless resistance management strategies are designed by elucidating the evolutionary and molecular basis of resistance, recent gains in reducing malaria burden could be lost [3, 4].

*Anopheles funestus,* a major malaria vector across sub-Saharan Africa, has demonstrated resistance to several insecticides, including pyrethroids, jeopardising recent efforts to eradicate and control malaria [5–19]. The primary mechanisms of insecticide resistance are target site resistance and metabolic resistance [20]. In *An. funestus,* metabolic resistance plays a predominant role, as the high expression of efficient detoxifying enzymes allows for the rapid removal or destruction of insecticides [21]. Metabolic resistance involves three-phase metabolic pathways present in all major groups of organisms. These pathways consist of modification, biotransformation, and excretion of toxic insecticides [21]. During Phase I, the toxicity of insecticides is reduced when a reactive and polar group is added to the substrate by a variety of enzymes including, cytochrome P450 monooxygenases (P450s), esterases, alcohol dehydrogenases and aldehyde dehydrogenases. In Phase II, the activated metabolites produced by the Phase I reactions are conjugated with charged species and bio-transformed into more polar and soluble metabolites that can be actively transported [22, 23]. The Phase II reactions are catalysed by a variety of transferases, including sulfotransferases, glutathione S-transferases (GSTs) and UDP-glycosyltransferases (UGTs) [24]. Conjugated toxins are then excreted from the cells into the extracellular medium via Phase III, where a variety of membrane-bound transporters are involved, notably ATP-binding cassette (ABC) transporters [24].

Previous studies have highlighted the role of *An. funestus* P450s enzymes, including CYP6P9a, CYP6P9b, CYP9J11, CYP6Z1, CYP6M7 and CYP9K1, in metabolising pyrethroids, leading to significant depletion and reduced efficacy of pyrethroids-treated bed nets [7, 8, 14, 17, 25, 26]. Additional classes of detoxification enzymes in *An. funestus*, such as GSTe2, confer resistance to pyrethroids and DDT through allelic variations that increase metabolic activities, resulting in cross-resistance [15]. The upregulation of other detoxification enzymes belonging to Phase II such as UGTs has been observed in previous transcriptomic studies investigating the molecular basis of resistance to pyrethroid [17, 26]. However, despite their established role in detoxification in other insects, UGTs role in pyrethroid resistance in malaria vectors, including *An. funestus*, remains largely uncharacterised.

UDP-glycosyltransferases (UGTs) constitute a superfamily of enzymes that play a vital role in the biotransformation of various hydrophobic compounds into more hydrophilic products [27]. These enzymes catalyse the covalent addition of a glycosyl group from an active donor, uridine diphosphate (UDP) glucose, to hydrophobic compounds containing hydroxyl, carboxyl, or amino functional groups through the glycosylation reaction [22, 27–29]. The resulting glycosides are more polar metabolites that can be easily excreted from the cell by export transporters than the substrate compound. Glucose conjugation is involved in various physiological processes in insects, including pigmentation, cuticle formation (sclerotization) and metabolic detoxification [30–32]. In certain lepidopteran insects, especially, UGT-mediated glycosylation activities are associated with resistance to plant defensive allelochemicals. These activities have been observed in economically important species such as the silkworm *Bombyx mori* [33], the tobacco hornworms *Manduca sexta* [34] and the Asian corn borer *Ostrinia furnacalis* [35–37] UGT-mediated biotransformation of pyrethroids has been suggested in *Anopheles sinensis,* a common malaria vector in Southeast Asia [38]. Their contribution to resistance against several classes of insecticide has been reported in the Diamondback moth *Plutella xylostella* and the tomato leafminer *Tuta absoluta* resistance to chlorantraniliprole [39, 40], the housefly *Musca domestica* resistance to organophosphate [41], and the greenfly cotton aphid *Aphis gossypii* resistance to neonicotinoid and spirotetramat [42, 43].

While genome-wide characterisation of UGT genes, their evolution, and association with insecticide resistance has been conducted in various insects using different sequencing technologies [44–47], UGT genes remain largely uncharacterised in malaria vectors. Although their overexpression in response to pyrethroids exposure has been reported in *An. funestus*, the investigations of UGTs in this context has been limited compared to other gene families, such as cytochrome p450s or GSTs. Selective sweeps and nonsynonymous mutations associated with UGTs, which could potentially contribute to pyrethroid resistance, have not been previously identified. In this study, we combine genome-wide association of pooled-template sequencing (GWAS-PoolSeq) with transcriptomic analysis of pyrethroids-resistant filed populations of *An. funestus* and deep sequencing of part of the genome of 80 mosquitoes to comprehensively characterise *An. funestus* UGTs genome-wide, their evolution, and association with pyrethroid resistance. This study provides the first comprehensive investigation into the role of UGTs role in pyrethroids resistance in *An. funestus*, the major malaria vector.

## 2. Methods

### 2.1. Identification of UGT genes, amino acids alignments, phylogenetic analysis and haplotype networks

*An. funestus* UGT genes were identified from the recently assembled and annotated *An. funestus* genomes based on protein family ID (Pfam ID) PF00201 from the recent protein family database Pfam 34.0 [48–51]. *An. funestus* UGTs amino acid sequences were retrieved from the *An. funestus* genome annotated protein and then were aligned in Geneious Prime 2022.1.1 (https://www.geneious.com) using Geneious alignment [51]. The signal peptide domain was predicted using SignalP 5.0 server (https://services.healthtech.dtu.dk/service.php?SignalP-5.0) and transmembrane helices were predicted in the amino acid consensus sequence using TMHMM - 2.0 (https://services.healthtech.dtu.dk/service.php?TMHMM-2.0) [52]. While UGTs conservative motifs and functional domains were identified by alignment with other insect UGTs amino acid sequences and using InterPro scan http://www.ebi.ac.uk/interpro/search/sequence/ [38, 44, 45].

For the phylogenetic analysis, amino acid sequences of 123 UGT genes from four Diptera species, *Anopheles funestus* (27), *Aedes aegypti* (35), *Anopheles gambiae* (26), *Drosophila melanogaster* (35) were retrieved from Vectorbase and globally aligned using Clustal Omega alignment built within Geneious Prime 2022.1.1 (https://www.geneious.com) [50]. The phylogenetic tree was built using Geneious built-in FastTree. FastTree uses a modified version of the neighbour-joining algorithm to construct an initial tree that is refined using a maximum-likelihood approach and heuristics are used to speed up the tree-building process [53]. The phylogenetic tree was edited using the online tool, Interactive Tree of Life (http://itol.embl.de/). *An. funestus* UGT genes were named based on the UGT nomenclature system [54].

To construct the Templeton, Crandall and Sing (TCS) haplotype network, UGT haplotypes for all individuals were extracted from the variant calling files using bcftools [55] resulting in 160 haplotypes for each UGT gene. Haplotypes were aligned using MUSCLE aligner [56] and TCS haplotype network was built using POPART software [57].

### 2.2. Mosquito collection, rearing and sequencing

The collection, rearing and sequencing of mosquitoes were described in detail previously in [7, 58]. In brief, two *An. funestus* laboratory colonies (FANG and FUMOZ) and field mosquitoes from Malawi, Cameroon and Uganda were used in this study. The FUMOZ colony is a multi-insecticide-resistant *An. funestus* colony derived from Southern Mozambique [59]. While the FANG is an insecticide-susceptible *An. funestus* colony derived from Angola [59]. Mosquitoes were collected from field populations representing southern Africa and Central Africa. Mosquitoes were collected from southern Chikwawa (16°1′S, 34°47′E), Malawi in January 2014 [13]; Tororo (0°45′N, 34°5′E), Uganda in March 2014 [12], and from Mibellon (6°46′N, 11°70′E), Cameroon in February 2015. Collected F_0_ gravid mosquitoes were forced to lay eggs using the forced egg-laying method [9]. The collected F_0_ mosquitoes were determined to be belonging to the *An. funestus* group using morphological and molecular identification [60, 61]. Egg batches were transported to the Liverpool School of Tropical Medicine (DEFRA) license (PATH/125/2012). Eggs were allowed to hatch, and mosquitoes were reared to adulthood in the insectaries under conditions described previously [9]. F1 females that are two-to-five old were subjected to insecticide resistance bioassays as described previously in [12, 13]. F1 females were exposed to permethrin for a varying length of time to define putatively resistant and susceptible mosquitoes. In populations from Malawi and Uganda, susceptible mosquitoes were defined as those that die after 60 min of permethrin exposure and resistant mosquitoes that are still alive after 180 min. In the population from Cameroon, due to lower levels of resistance susceptible mosquitoes were dead mosquitoes collected after 20 minutes of exposure and resistant are those that are still alive after 60 min exposure. Genomic DNA was extracted using DNeasy blood and tissue kit (Qiagen) from 40 mosquitoes individually and pooled in equal amounts. Library preparation and whole-genome sequencing by Illumina HiSeq2500 (2 × 150 bp paired-end) were carried out by the Centre for Genomic Research (CGR), University of Liverpool, UK [17, 18].

For transcriptional profiling of pyrethroid-resistant in field-collected populations, a total of 18 RNA pools were collected for populations from Malawi, Cameroon and Uganda. In each field-collected population, 3 pools of alive mosquitoes after exposure to permethrin and 3 pools of unexposed mosquitoes were collected. In addition, 4 pools each were collected for laboratory colonies FANG and FUMOZ. Total RNA was extracted from pools of 10 female mosquitoes using the Arcturus PicoPure RNA Isolation Kit (Life Technologies) as described previously [17, 26]. Library preparation and sequencing by Illumina HiSeq 2500 (2 x 125-bp) were done at the Centre for Genomic Research (CGR), University of Liverpool [17, 18].

### 2.3. Design of the SureSelect bait and target enrichment sequencing of candidate resistance regions from laboratory and field-collected colonies

Target sequencing baits were designed using the SureSelect DNA Advanced Design Wizard in the eArray program of Agilent. The design and sequencing of the SureSelect experiment were described in detail previously [7] In summary, a total of 1302 target sequences were included in the enrichment sequencing, constituting 3,059,528 bp. The library preparation and sequencing were performed by the Centre for Genomic Research (CGR), University of Liverpool, using the SureSelect target enrichment custom kit. The libraries were sequenced in 2 × 150 bp paired-end fragments on an Illumina MiSeq with 20 samples per run. The regions targeted by the enrichment sequencing include a selection of detoxification genes potentially involved in insecticide resistance, heat shock proteins, immune response genes and odorant binding proteins (see [7] for further details). In addition, all the genes in the major candidate trait loci associated with pyrethroid resistance previously identified were included in the targeted sequencing. These include a 120 kb region Bacterial Artificial Chromosome (BAC) clone of rp1 (resistance to pyrethroid 1) locus on chromosome 2R and a 113 kb BCA clone of rp2 on chromosome 2L [19, 62].

This fine-scale targeted technique was used to sequence part of the genome of a total of 80 individual mosquitoes from the two lab colonies (FANG and FUMOZ) and filed-collected mosquitoes from Cameroon, Malawi and Uganda. For field-collected populations, the set used for PoolSeq, from Malawi, Uganda and Cameroon,10 putatively permethrin susceptible and 10 resistant mosquitoes were targeted by the SureSelect bait. In addition, 10 mosquitoes were targeted from each laboratory colony FANG, FUMOZ. Centre for Genomic Research (CGR), the University of Liverpool conducted the library construction and capture using the SureSelect target enrichment custom kit. The libraries were subjected to paired-end sequencing (2 × 150 bp) on an Illumina MiSeq instrument with 20 samples per run [7]. A broad-scale analysis of these data can be found in [7].

### 2.4. Differential expression analysis of total RNA and analysis of PoolSeq data

Differentially expressed genes (DEGs) analyses were conducted between the laboratory colonies FANG and FUMOZ, between putatively susceptible and resistant populations in each filed-collected population, and between resistant filed-collected populations and the FANG colony. The analysis involves initial pre-processing of raw reads, alignment to the reference genome using the AfunF3.2 annotation and count of reads on the gene level. The DEG analysis was conducted using edgeR [63]. Differentially regulated genes were defined in each separate contrast as those with a corrected p-value threshold of < 0.05 and log_2_ fold change > 1.

To validate the expression profiles of select UGTs using quantitative real-time PCR, RNA was extracted in pools of 10 from 3-5 days old females randomly selected and unexposed to any insecticides using Arcturus PicoPure RNA isolation kit (Applied Biosystems, CA, USA) according to the manufacturer’s instructions. Reverse transcription and qRT-PCR were preformed using Invitrogen reverse transcriptase kit (Invitrogen, Waltham, CA, USA) and SYBR green master mix kit (Sigma, Aldrich, Germany) respectively according to the manufacturer’s instructions. The expression of RNA was calculated using ddCT protocol [64] and normalised to the expression of Ribosomal protein S7 housekeeping gene. The efficiency of the primers was incorporated into the analysis and values were finally converted into their logarithmic forms for normal distribution. The primer sequences are provided in Supplementary Table S9.

For the PoolSeq, the pre-processing of raw reads, alignment and variant calling was described previously [7]. Gene-wise pN/pS was calculated using snpgenie [65] for details see [58].

### 2.5. Analysis of the SureSelect data

SureSelect pair-end trimmed reads were aligned using BWA (version 0.7.17) against *An. funestus* reference genome [51]. Sequence alignment map (sam) files were converted to binary alignment map (bam) files using samtools (1.13) [66, 67]. Using Picard tools (2.26.3) alignment bam files were sorted according to the position in the reference genome, duplicated reads were marked, and reads were assigned a new read group tag [68]. Variant calling was carried out using freebayes (v1.2.0-dirty) by setting the number of alleles to be considered to 4 using the option (--use-best-n-alleles 4) to reduce run time and all other options were set to default [69]. Subsequently, resulting variants were filtered using vcffilter (vcflib version 1.0.0_rc2) based on Phred-scaled quality-score greater than 20 (QUAL >= 20) keeping 531726 variants [70]. SnpEff (5.0) was used to annotate and predict the effects of genetic variants between samples and the FUMOZ reference genome [71]. Variants were further filtered by removing SNPs with missing values, removing all indels, and only retaining bi-allelic SNPs. Variants were separated by country to multiple VCF files so when filtered by missing values a maximum number of SNPs will be retained (Supplementary Table S6) [66, 67].

### 2.6. Gene-wise *F_ST_* differentiation and allele frequency spectrum (AFS) test statistics between pyrethroid resistance and putatively susceptible populations

To conduct population genetics analyses on a gene level from the SureSelect target enrichment sequencing only genes with an average coverage of 100 and above calculated in jvarkit https://github.com/lindenb/jvarkit using merged alignment files of all the FANG and FUMOZ individuals were retained. The target enrichment sequencing covered 807 genes, compromising 6.16% of the total number of genes in the FUMOZ reference genome. Gene-wise *F_ST_* association between resistance and susceptible mosquitoes within each country, Africa-wide, between lab colonies (FANG and FUMOZ) was calculated using PopGenome R package [72]. Gene-wise population genetics neutrality statistics including Tajima’s D (on synonymous SNPs, nonsynonymous SNPs and combined) Fu, and Li‘s D and F tests, Watterson estimator and composite likelihood ratio (CLR) test [73] were calculated for all genes included by the targeted sequencing using PopGenome R package [72]. Coalescent simulation to derive the expected distribution of gene-wise neutrality tests statistics across the loci targeted by the enrichment analysis was generated using the MS program [74].

### 2.7. *F_st_* differentiation between pyrethroid resistance and putatively susceptible populations on the SNP level

For all SNPs in the targeted region by SureSelect, Weir & Cockerham’s *wc F_ST_* for two populations and *p F_ST_* probabilistic approach to detect the difference in allele frequencies between the resistant and susceptible population in every country, Africa-wide and between laboratory colonies (FANG and FUMOZ) was calculated using vcflib (1.0.0_rc2) [70]. R qvalue package (https://github.com/StoreyLab/qvalue) was used to perform a false discovery rate (FDR) estimation for the list of p-values calculated from the *p F_ST_* test, a p-value of 0.05 were used to extract significantly divergent SNPs from each analysis. The functional effect of each SNP with significant *p F_ST_* from each comparison on the coding region was determined by overlapping using SNPs genomic positions. Supplementary Table S6 surmises the total number of SNPs identified in the targeted region, along with the significantly divergent SNPs between resistant and susceptible, and the number of predicted functional effects of significant SNPs including non-synonymous SNPs. The functional effect of each SNP as predicted by SnpEff, a p-value quantifying allele frequency difference (*p F_ST_*) and Weir & Cockerham’s *F_ST_* (*wc F_ST_*) are provided in (Supplementary Dataset 7).

## 3. Results

### 3.1. Characteristics, phylogenetics, and evolution of *An. funestus* UDP-glycosyltransferases (UGT) genes

The assembled and annotated *Anopheles funestus* genome for the FUMOZ resistant-strain colony contains 21 UGTs; 20 genes with complete transcripts located on chromosomes 2 and 3, encoding 511 to 550 amino acids (Table 1), and a partially sequenced UGT gene AFUN018708 (65 amino acids) due to incompleteness in the transcriptome assembly [51]. The latest *An. funestus* assembly from an individual female specimen from La Lopé, Gabon (idAnoFuneDA-416_04) contains seven more UGT genes and the partially sequenced UGT gene (AFUN018708) from the FUMOZ colony reference was identified to be a partial sequence of AFUN2_007305(b) later named *UGT308B3* according to UGT nomenclature [75]. Overall, the number of predicted genes in the idAnoFuneDA-416_04 assembly (14,819 genes) is higher than the FUMOZ colony assembly (14,176 genes). In total 27 UGT genes were identified in *An. funestus* from both assemblies. (Table 1) [51]. Genomics and transcriptomics analyses in this paper used the FUMOZ strain reference genome as its annotation is well-integrated within VectorBase.

**Table 1.**
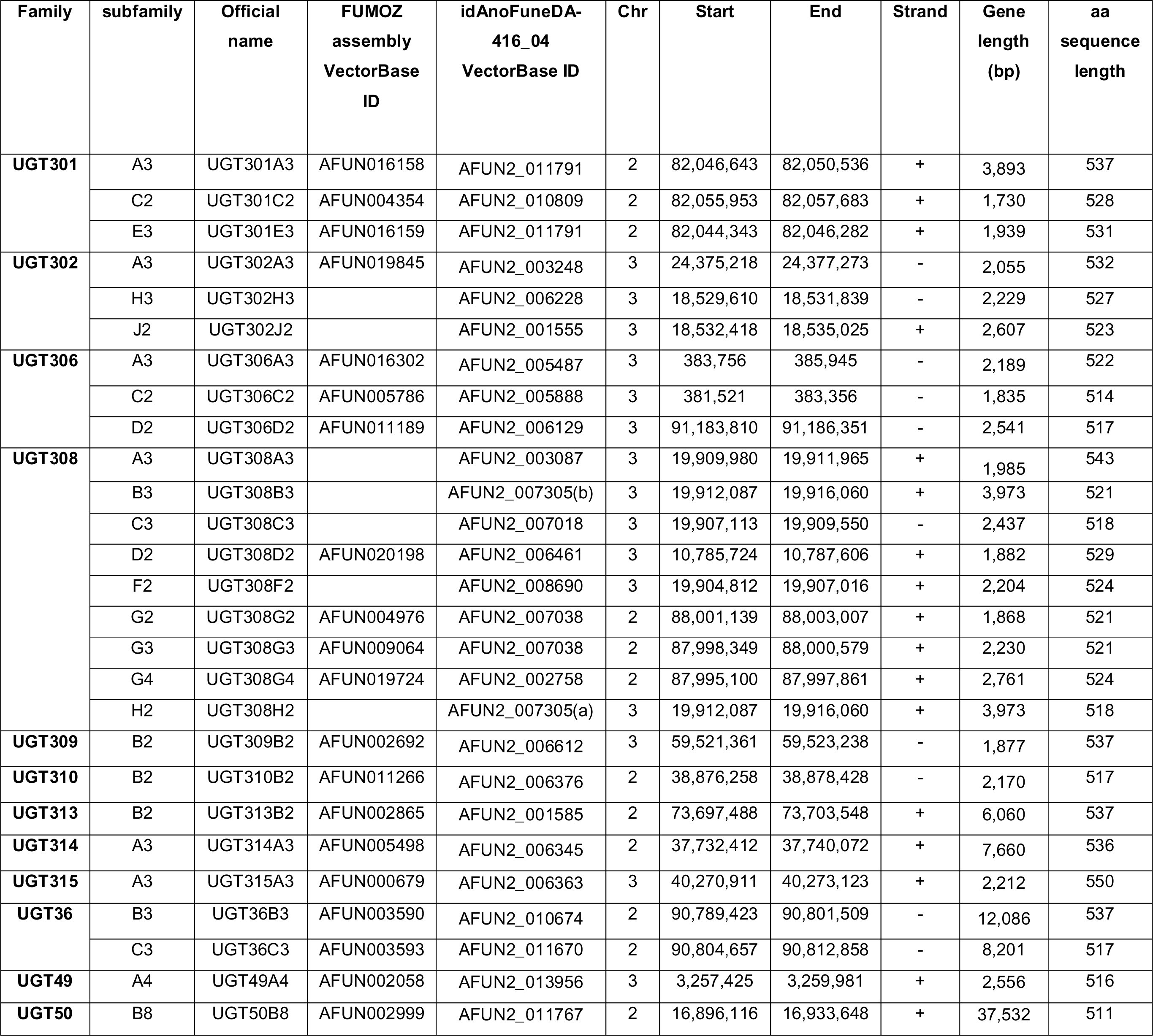
A complete list of UGT genes identified in *An. funestus*. The table includes official names for all UGT genes, VectorBase gene IDs, chromosomal location including gene start (Start), end position (End) and strand orientation (Strand), and length of the gene and amino acid sequence (aa sequence length). For the overlapping 20 UGT genes found in the FUMOZ colony reference and in the idAnoFuneDA-416_04 genome, VectorBase IDs from both genomes were reported, while Chr, Start, End and Strand were extracted from the FUMOZ reference genome. For the extra 7 UGTs specific to idAnoFuneDA-416_04 genome, the chromosome location, gene start, and end position were reported from the idAnoFuneDA-416_04 genome.

Phylogenetic analysis of a total of 123 protein sequences of UGT genes from four Diptera species, *Anopheles funestus* (27), *Aedes aegypti* (35), *Anopheles gambiae* (26), and *Drosophila melanogaster* (35), divides UGT genes to 20 canonical families according to the UGT nomenclature committee (Fig. 1) (Supplementary Dataset 1) [54]. The phylogenetic tree reveals lineage-specific expansion and interspecific conservation of the UGT families. Five out of the 20 families are common in all four Diptera species (UGT36, UGT49, UGT50, UGT301 and UGT302), while 8 families are specific to *Drosophila melanogaster* (UGT35, UGT37, UGT303, UGT304, UGT305, UGT307, UGT316, and UGT317) and the remaining 7 families are specific to Culicidae species (UGT306, UGT308, UGT309, UGT310, UGT313, UGT314 and UGT315).

**Fig. 1.**
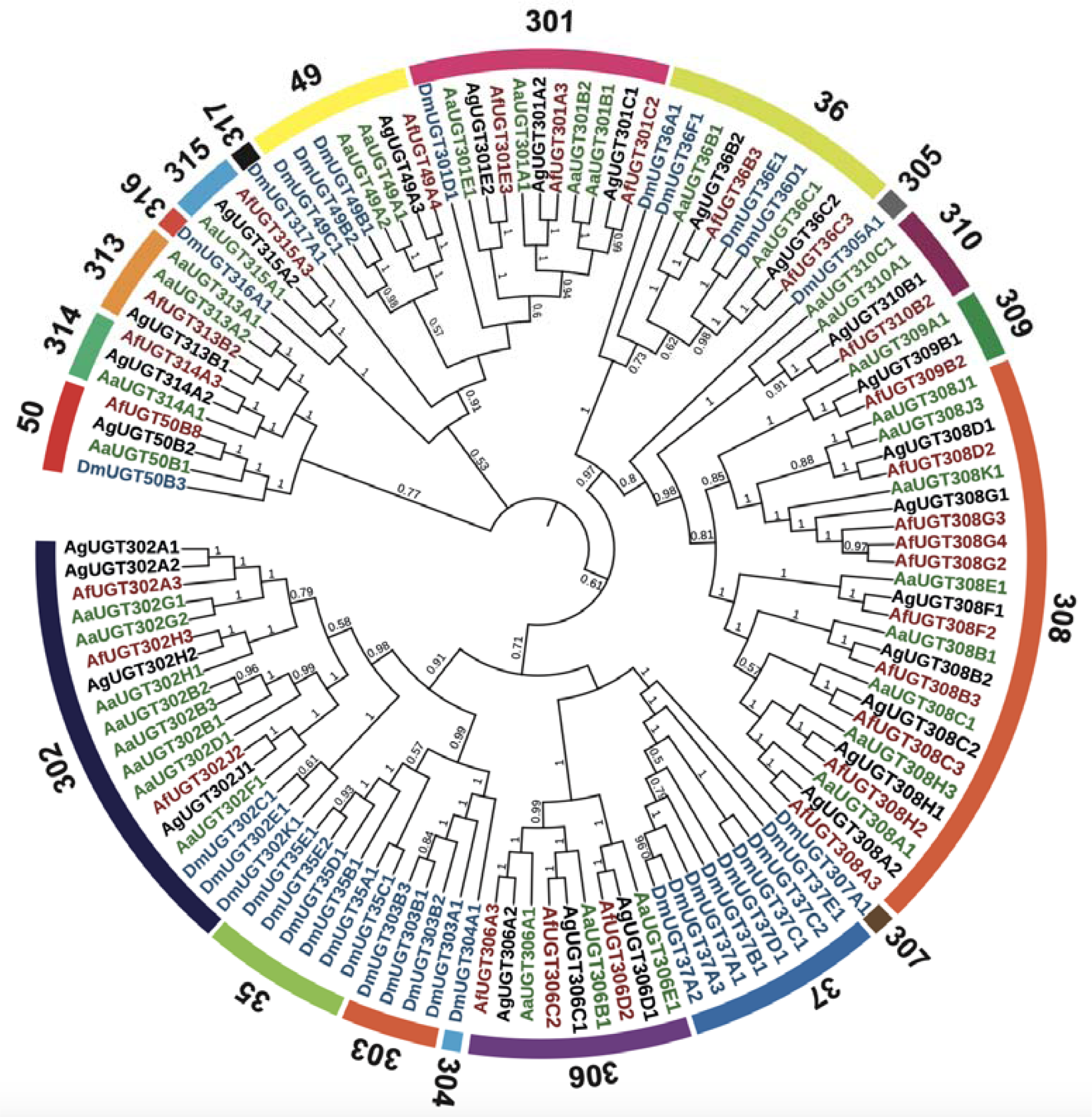
Phylogenetic tree of 123 UGT sequences from *An. funestus*, *Ae. Aegypti*, *An. gambiae*, *D. melanogaster*. Phylogenetic analysis was constructed using FastTree method. Fast tree support values that are larger than 0.5 are marked on the corresponding branch. UGT names start with the species initials *An. funestus* (Af), *Ae. Aegypti* (Aa), *An. gambiae* (Ag), and *D. melanogaster* (Dm), followed by the family classification (numbers), the subfamily classification (letters), and the unique numbers. The numbers on the outer circle refer to the family classification.

*Anopheles funestus* UGT genes were distributed into 12 families, of which 9 UGTs were clustered within the UGT308 family (Fig. 1, Table 1). The UGT308 gene family demonstrates significant gene expansion, making it the largest in the overall phylogenetic tree and the largest among the UGT families in *An. funestus*. There are 3 *An. funestus* UGT genes each in UGT families UGT301, UGT302 and UGT306 with a close 1:1 orthologue gene between *An. funestus* and *An. gambiae*. There are two *An. funestus* UGT genes in the UGT36 family belonging to subfamilies B and C are close orthologous to the UGTs from *An. gambiae* and *Ae. Aegypti* belonging to the same subfamilies. There is only one *An. funestus* UGT each in UGT49, UGT50, UGT309, UGT310, UGT313, UGT314 and UGT315 families (Fig. 1, Table 1). The species phylogeny among the four Diptera species is mirrored by the branching pattern of the UGT50 family, the *An. funestus* UGT50 gene is closer to *An. gambiae* and *Ae. Aegypti* with a pairwise identity of 93.1% aaID and 79.6% aaID respectively than *D. melanogaster* DmUgt50B3 with 58.6% aaID (Fig.1) (Supplementary Dataset 2).

Protein function analysis using the consensus sequence and UGT50B8 amino acid sequence predicted a signal peptide at the N-terminal, a transmembrane domain, and a cytoplasmic tail at the C-terminal domain (Supplementary Fig. 2). A significant similarity is observed at the UGT signature motif of 29 aa-long sequences (consensus: FITHGGLLSTQEAIYHGVPIVGIPFFGDQ) found in the middle of the C-terminal domain with two residues conserved in all UGT genes (G415, P428) [54]. Other conserved signature motif sequences involved in sugar donor binding regions (DBR1 and DBR2) and catalytic mechanisms are found in the C-terminal domain comparable to mammalian and insect UGT genes (Supplementary Fig. 1).

### 3.2. Studying UGT association with pyrethroid resistance using DNA and RNA pools

All the 21 UGT genes annotated in the FUMOZ genome including the partially sequenced UGT308B3 (AFUN018708) were investigated for association with pyrethroid resistance. DNA pools (PoolSeq) from *An. funestus* laboratory colonies (FANG and FUMOZ) and field-collected mosquitoes from Malawi, Uganda and Cameroon were examined for a signature of selection in the UGT gene family [7, 58]. In each population, Pools of genomic DNA were analysed by calculating the gene-wise ratio between nucleotide diversity of nonsynonymous polymorphism and nucleotide diversity of synonymous polymorphism (π_n_/π_s_) for all genes including the 21 UGT genes (Supplementary Dataset 3) [7, 58]. The (π_n_/π_s_) ratio gene-wise for all the UGT genes in all the populations is generally less than 1, implying stabilizing or purifying selection acting against changes in the amino acid sequence, except for *UGT315A3* (AFUN000679) in the FUMOZ population where a positive selection was detected driving changes in the protein sequence.

To identify UGTs that are overexpressed in response to exposure to permethrin, differentially expressed genes (DEGs) analyses between mosquitoes alive after exposure to permethrin and unexposed populations from Malawi, Uganda, and Cameroon were performed using pools of total RNA. In addition, differentially expressed gene analyses were performed by contrasting transcription profiles of populations resistant to permethrin from Malawi, Uganda, Cameroon, and laboratory-resistant colony (FUMOZ) against the transcription profile of laboratory-susceptible (FANG) (See Materials and Methods for details). In each separate contrast, DEGs were determined globally while DEGs of detoxification genes were highlighted including differential expressions of UGT genes (Supplementary Fig. 3). Differential expression analyses between mosquitoes resistant and unexposed to permethrin in populations from Uganda and Cameroon did not detect significant changes in the expression level of detoxification genes including UGTs. However, the expression contrast in Malawi between resistant and unexposed populations detected significant differential expression of 7 UGT genes; 5 genes were upregulated, and 2 genes were downregulated. In Malawi, 4 of the genes that are upregulated are also upregulated when the resistant population is compared with FANG (Fig. 2). *UGT310B2* (AFUN011266) is significantly upregulated in all resistant populations to permethrin when compared to the FANG population, 2.7 FC in FUMOZ, 5.6 in Malawi, 2.6 in Uganda and 3.5 FC in Cameroon (Fig. 2B). While *UGT301C2* (AFUN004354) overexpression was specific to the FUMOZ population (2.2 FC) when compared with FANG (Fig. 2, Supplementary Fig. 4, Supplementary Dataset 4). The overexpression of *UGT310B2* in FUMOZ compared to FANG was detected using quantitative PCR (qPCR) (Fig. 2C). Detecting *UGT301C2* by qPCR was challenging due to its relative overexpression in the FUMOZ colony compared to the FANG (Fig. 2C, Supplementary Dataset 4).

**Fig. 2.**
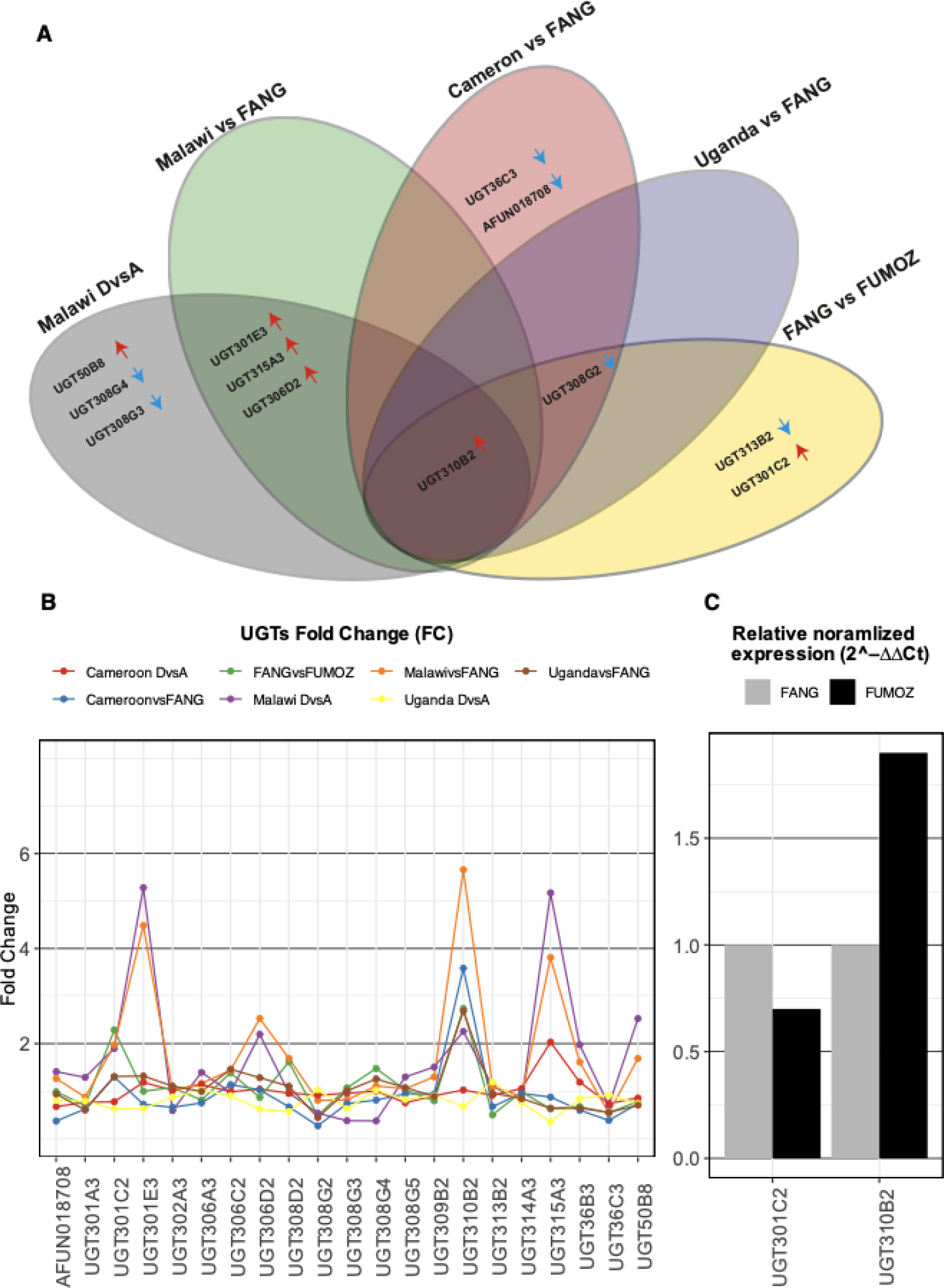
Differentially expressed *An. funestus* UGT genes in response to permethrin resistance. **(A)** Differentially expressed genes (DEGs) between the field-resistant population after exposure to permethrin and the unexposed population from the same country were identified in Malawi but not in Cameroon and Uganda. **(A)** Comparing the transcriptional profile of resistant populations from Malawi, Cameroon, Uganda and FUMOZ to that of susceptible FANG population identified Africa-wide upregulation of *UGT310B2*. The red arrow pointing upwards indicates upregulation and the blue arrow pointing downwards indicates downregulation. **(B)** All *An. funestus* UGTs (x-axes) plotted against fold change (FC) (y-axes) obtained from DEGs analyses colour coded according to the plot legend highlights the Africa-wide upregulation of *UGT310B2* with a maximum fold change of 5.6 in resistant population from Malawi when compared with transcription profile of FANG. *UGT310B2, UGT306D2, UGT301E3,* and *UGT315A3* have a significant overexpression in Malawi resistant population when compared with the FANG profile and with the unexposed population from Malawi highlighting the potential role of UGTs in detoxification in Malawian population. **(C)** The overexpression of *UGT310B2* was detected in the FUMOZ colony using qPCR. However, due to the relative overexpression of *UGT301C2* from the FUMOZ RNAseq profile compared to the FANG detecting the overexpression in the FUMOZ colony by qPCR was challenging.

Overall, global transcriptional regulation in resistant populations in response to permethrin exposure, when compared to unexposed populations from the same region, was more evident in Malawi compared to Uganda and Cameroon (Supplementary Fig. 5). In Malawi, a total of 1826 significant DEGs were detected, and 48 of those genes belong to the major detoxification gene families, 21 cytochrome P450s, 12 carboxylesterases, 4 glutathione S-transferases, 4 UDP-glycosyltransferases and 2 ABC transporters. In the population from Cameroon, only a single carboxylesterase gene (AFUN002514) was detected to be upregulated [17]. While the Ugandan resistant population overexpresses three cytochromes P450 genes AFUN015739 (*CYP307A1*), AFUN020895 and AFUN019365 compared to the unexposed population from the same country (Supplementary Fig. 3, Supplementary Dataset 4) (Supplementary text 1).

### 3.3. Identifying selection and divergence in *An. funestus* UGT genes using targeted sequencing

UGT genes associated with pyrethroid resistance were detected using gene-wise allele frequency spectrum (AFS)-based methods from a targeted and fine-scale sequencing technique used to enrich part of the genome of 80 individual mosquitoes (see methods). Biallelic SNPs within the targeted regions for each individual mosquito were used in downstream gene-wise allele frequency spectrum-based (AFS) summary statistics. A total of 136,348 bi-allelic polymorphic sites for AFS analysis were retained across 80 mosquitoes Africa-wide, divided into equal numbers of dead and alive mosquitoes across the continent and in each country (see methods). In each country, 91,619, 90,340, and 49,762 polymorphic biallelic sites were detected respectively in Malawi, Uganda and Cameroon (Table S1). polymorphic sites were divided between genes targeted using the enrichment sequencing that includes 61 genes on the X chromosome, 431 genes on Chr2 and 315 genes on Chr3 (807 in total) (Table S1). Many of those genes belong to detoxification gene families, including P450 (66 genes), GSTs (12 genes), COEs (5 genes), UGTs (12 genes), ABC transporters (14 genes), and some of the remaining 698 genes could also be associated with pyrethroid resistance (Supplementary Text 2).

Low polymorphism was detected in samples collected from Malawi compared to populations from other countries. Additionally, low polymorphism was detected between laboratory colonies FANG and FUMOZ compared to all filed isolates, where 28,566 polymorphic sites were detected. A low level of polymorphism in the field population from Malawi and the FANG laboratory colony, originally from Angola, is expected when FUMOZ, originally from Mozambique, is used as a reference genome since all populations are from southern Africa (Table S1).

For each gene targeted by the enrichment sequencing, gene-wise *F_ST_ was* calculated to investigate divergence between resistant (alive) and susceptible (dead) Africa-wide, in each country and between laboratory colonies (FANG and FUMOZ). Genes with gene-wise *F_ST_* values that are in the top 0.05 quantiles of genes per chromosome were considered significant (Supplementary Dataset 5, Fig. 3, Supplementary Fig. 6). To detect genomic regions impacted by selections, gene-wise Tajima’s D neutrality test statistics were estimated for all genes included in the targeted sequencing. The significance of gene-wise Tajima’s D values of all genes targeted by the enrichment sequencing including UGT genes were determined from Tajima’s D density plot of the coalescence simulation (See Materials and Methods for details). Tajima’s D values were considered significant if they fall at the 0.05 quantiles at both extremities of the simulated Tajima’s D density plot (Fig. 4, Supplementary Dataset 6).

**Fig. 3.**
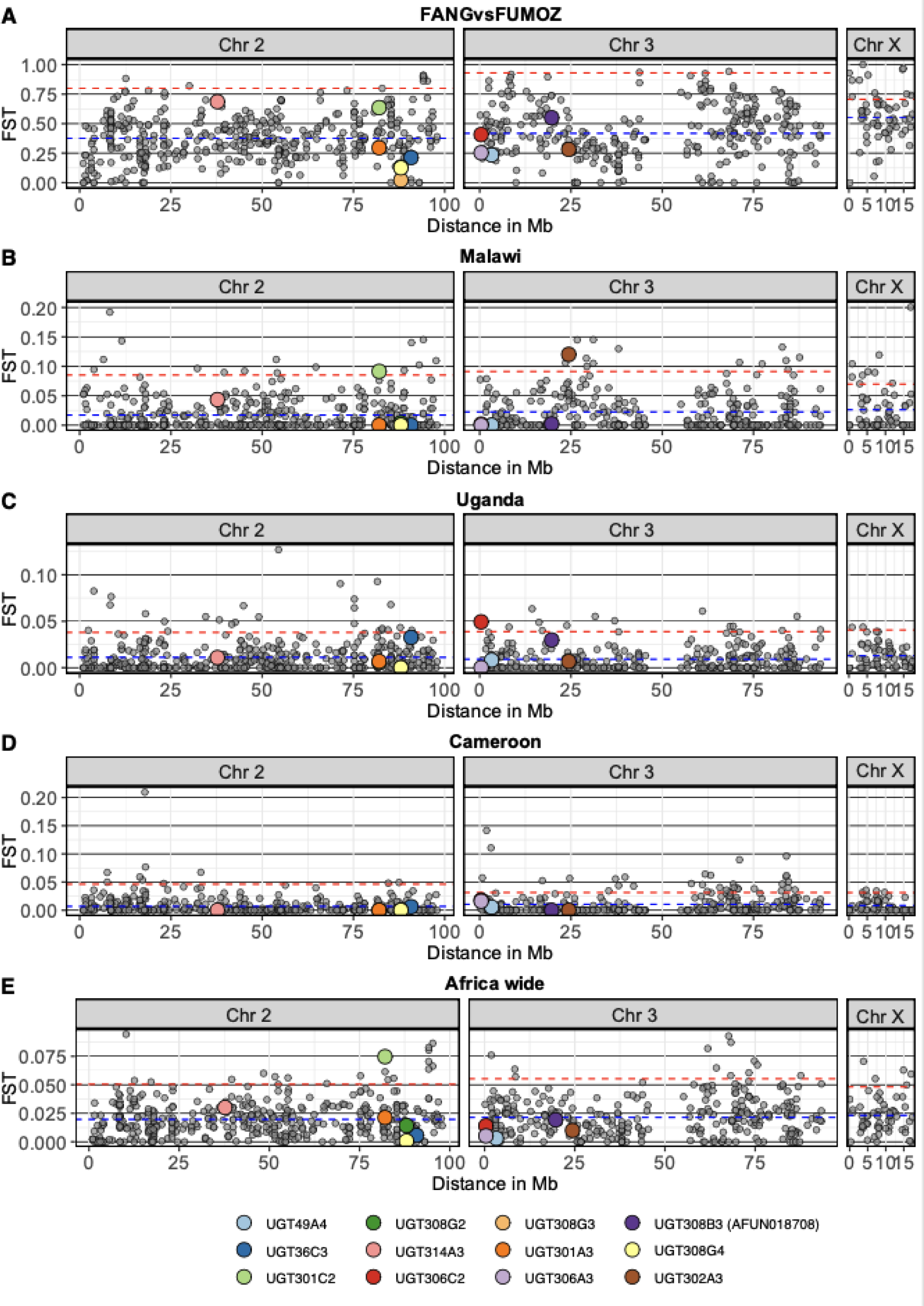
Gene-wise *F_ST_* for all genes included in the targeted sequencing in all analyses between resistant and susceptible. The average and 0.95 quantiles of gene-wise *F_ST_* for each chromosome are represented by the blue horizontal dotted line and the red dotted line respectively. The genomic location of all genes in the targeted region (x-axis), represented by a circle, was plotted against the Gene-wise *F_ST_* (y-axis). UGT genes included in the targeted sequencing are colour coded according to the plot legend. Raw data of all the genes included in the targeted sequencing is in (Supplementary Dataset 5), averages and 0.9 quantiles for each comparison are in (Table S2), and UGT genes *F_ST_* values are in (Table S3).

**Fig. 4.**
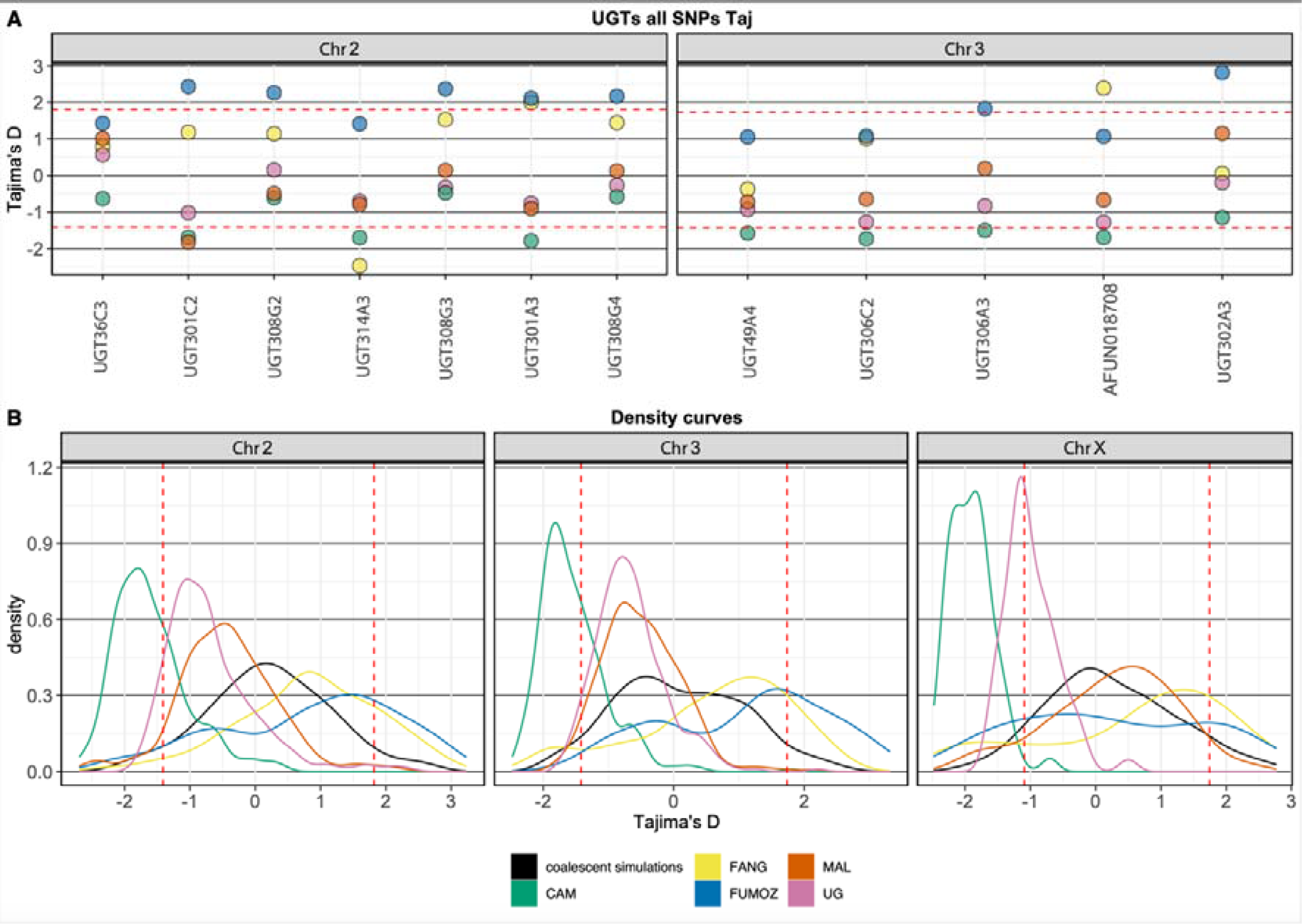
Tajima’s D for UGT genes and density curves of all the genes included by the targeted enrichment sequencing for all the populations. In populations from southern Africa Malawi (orange), FANG (Angola) (Yellow) and FUMOZ (Mozambique) (Blue) Tajima’s D density curves were close to equilibrium (Black) **(B)**. In Malawi, *UGT301C2* is impacted by a recent selective sweep with a negative below the 0.05 quantiles of simulated Tajima’s D **(A)**. In Uganda and Cameroon, gene-wise Tajima’s D was predominantly skewed towards negative values. In Cameroon, 7 UGT genes are below 0.05 quantiles of simulated Tajima’s D and most UGT genes in Uganda are with a negative Tajima’s D but not within the lowest 5% **(A)**.

Africa-wide gene-wise *F_ST_* test between dead and alive mosquitoes across the investigated countries detected a significant gene-wise *F_ST_* value of 0.07463 for UGT301C2 on the top 0.5 quantiles (Chromosome 2 0.95 quantiles= 0.048) (Fig. 3) (Table S2 and S3). To some extent, a geographical pattern of elevated gene-wise *F_ST_* for UGT genes was evident in samples collected from southern African populations (FANG, FUMOZ and Malawi) with elevated gene-wise *F_ST_* for *UGT301C2* and *UGT314A3*, compared to Uganda (central east) and Cameroon (central west) with a shared high gene-wise *F_ST_* for *UGT306C2* (Fig. 3) (Table S3). The commonly overexpressed *UGT310B2* was not included in the targeted sequencing to investigate if there is a link between differentiation and overexpression. *UGT301C2 which* is overexpressed in the FUMOZ colony is highly differentiated between FUMOZ and FANG populations but not significant. Gene-wise Tajima’s D density curve of genes included in the enrichment sequencing may reflect the demographic history of *An. funestus* population in those countries (Fig. 4 and Supplementary Fig. 7).

#### 3.3.1. Gene-wise differentiation and selection in laboratory colonies (FANG and the FUMOZ) and Malawi

Gene-wise *F_ST_* test statistics were employed to compare the susceptible laboratory colony the FANG, originally from Angola (southwest Africa), and the resistant laboratory colony the FUMOZ originally, from Mozambique (southeast Africa). No UGTs with significant elevated gene-wise *F_ST_* were detected amongst the top 0.05 quantiles in respective chromosomes, but *UGT301C2* and *UGT314A3* on chromosome 2 are the most differentiated UGT genes (Fig. 3) (Table S2). The average gene-wise *F_ST_* values in the analysis between FANG and FUMOZ were higher than all the other analyses between putatively susceptible and resistant from other countries, revealing the genetic difference between the two geographically distant isolates (Fig. 3) (Table S2). Furthermore, in Malawi (southeast Africa), significant differentiation of UGT genes *UGT301C2* on chromosome 2 and *UGT302A3* on chromosome 3 was identified, amongst the top 0.05 quantiles on respective chromosomes (0.95 quantiles of chromosome 2 = 0.069 and chromosome 3 = 0.085), with gene-wise *F_ST_* of 0.9142 and 0.12021 respectively. (Fig. 3 and Table S3). Meanwhile, differentiation of *UGT314A3* on chromosome 2 in populations from Malawi was similar to the observed differentiation between laboratory colonies (the FANG and the FUMOZ) only higher than 80% of gene-wise *F_ST_* values and not significant (Fig. 3).

In all populations from southern Africa including Malawi, FANG colony (derived from Angola) and FUMOZ colony (derived from Mozambique) Tajima’s D density curves were close to equilibrium, represented by the coalescent simulation curve (Fig. 4B). In the FUMOZ population, 7 UGT genes out of the 12 UGTs targeted in this study have a high gene-wise Tajima’s D than 0.95 quantiles of simulated Tajima’s D values and the gene-wise Tajima’s D average in FUMOZ is higher than the average of simulated Tajima’s D for all chromosomes (Fig. 4) (Table S4 and S5). Similarly, in the susceptible FANG population, Tajima’s D average for all chromosomes is higher than the coalescent simulation average (Table S4). This observation may indicate a strong balancing selection or decrease in population size (sudden population contraction) in the laboratory colony populations, probably introduced by laboratory propagation. However, in Malawi, *UGT301C2* has a negative Tajima’s D value of - 1.8243 below the 0.05 quantiles of simulated Tajima’s D, deviating from the empirical distribution of gene-wise Tajima’s D values on chromosome 2 indicating a recent selective sweep driven by directional selection (Fig. 4, Table S5).

#### 3.3.2. Gene-wise differentiation and selection in Central West Africa (Cameroon) and Central East Africa (Uganda)

When comparing susceptible and resistant populations from Uganda, UGT306C2 on chromosome 3 has a significant differentiation with gene-wise with *F_ST_* value of 0.0494 within the top 0.05 quantiles, while in Cameroon gene-wise *F_ST_* for *UGT306C2* is high but not significant (Fig. 3, Table S3 and Supplementary Dataset 5).

In Uganda and Cameroon, gene-wise Tajima’s D values were predominantly skewed towards negative values. In Cameroon, the average and median of gene-wise Tajima’s D per chromosome are below the 0.05 quantiles of simulated Tajima’s D for corresponding Chromosomes. There were 7 UGT genes in Cameroon below 0.05 quantiles of simulated Tajima’s D and most UGT genes in Uganda have negative Tajima’s D values but not within the lowest 5% of simulated Tajima’s D for corresponding chromosomes (Fig. 4, Supplementary Fig. 7, Table S4 and S5).

### 3.4. Analysis of UGTs polymorphism across Africa identified nonsynonymous SNPs potentially associated with resistance

The genomic fragment spanning the full length of UGTs targeted by enrichment sequencing were analysed in 80 mosquitoes Africa-wide. Nucleotide diversity (π) was calculated within each population (resistant and susceptible) as well as between populations from Malawi, Uganda, and Cameroon, and both laboratory colonies the FANG and the FUMOZ. Low gene-wise nucleotide diversity was detected in the susceptible FANG population for all UGT genes compared to other populations probably due to the lack of selection pressure and consistent population genetic drift introduced by laboratory maintenance of the colony (Fig. 5A). The low diversity of the FANG UGTs haplotype is evident in haplotype networks were FANG haplotypes mostly cluster together. Gene-wise nucleotide diversity varies between UGT genes, and the number of polymorphic substitutions relates to the gene size (Fig. B). Analysis of the Templeton, Crandall and Sing (TCS) haplotype tree for UGTs targeted by the enrichment sequencing from 80 mosquitos (160 haplotypes) highlights the high polymorphism of UGTs across Africa with many singleton haplotypes separated by many mutational steps, except for *UGT306C2*, *UGT306A3* and *UGT301C2* where predominant haplotypes shared by different populations were detected (Fig. 5C-E).

**Fig. 5.**
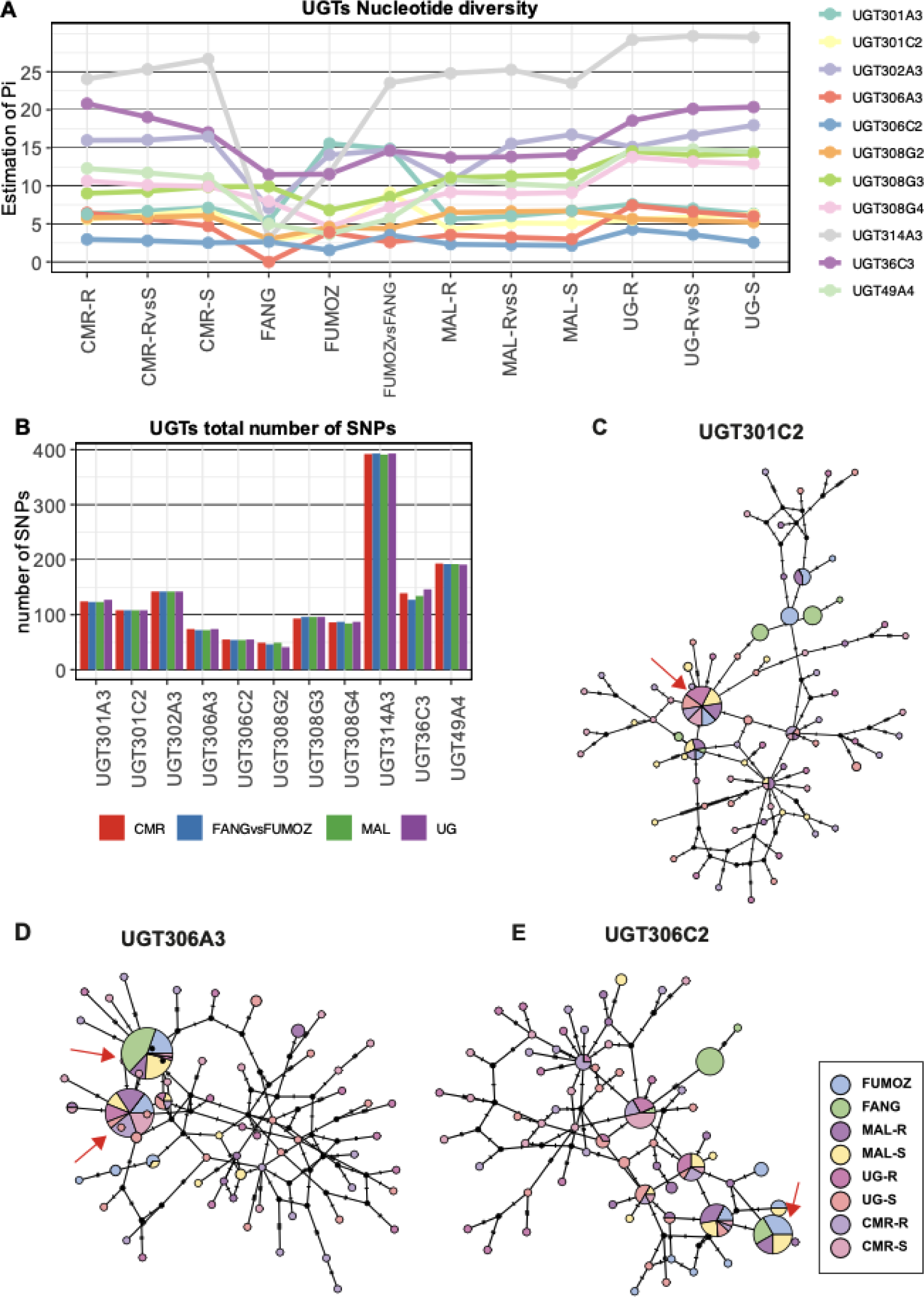
Targeted UGTs polymorphism and haplotype network analysis. UGTs gene-wise nucleotide diversity was calculated between (resistant and susceptible) populations and within populations from Malawi, Cameroon, Uganda and Laboratory colonies the FANG and the FUMOZ (A). The number of polymorphic substitutions for UGTs in each country reflects the gene size (Table 1) (B). TCS haplotype network reveals limited haplotype clustering in *UGT301C2* (C), *UGT306A3* (D) and UGT306C2 (E) where predominant haplotypes shared by different populations were detected. The predominant haplotype in UGT301C2 was observed in (40/140) haplotypes from Cameroon (9), Uganda (14), Malawi (12) and FUMOZ (5). While 18/20 FANG haplotypes cluster together in three predominant nodes separated by a single mutation step each contains (9/20), (7/20), and (2/20) haplotypes respectively (Fig. 5C). In *UGT306A3* two main haplotype clusters separated by a single mutation step were detected. The biggest cluster contains (45/160) haplotypes where all the 20 FANG haplotypes cluster, 9/20 FUMOZ and 15/40 haplotypes from Malawi. The second cluster contains (41/160) haplotypes including 12/40 from Malawi, 16/40 from Cameroon and 7/40 from Ugandan population (Fig. 5D). In *UGT306C2* many haplotype clusters were observed the most dominant is 23 haplotypes cluster specific to FUMOZ (7/20), FANG (6/20) and Malawi (11/40).

To identify SNPs that are potentially associated with pyrethroid resistance within UGTs, we identified significantly differentiated SNPs based on p-values quantifying differences in allele frequencies (*pFst*) between susceptible and resistant populations in each country (Table S6 and S7) In comparison between laboratory colonies FANG and the FUMOZ, 30.5% of targeted UGT SNPs show significant differentiation, including 26 non-synonymous variants. In Malawi, 11.6% of UGT SNPs are significantly differentiated, with 6 genes containing 14 significant nonsynonymous SNPs. Cameroon shows 5.4% of significantly differentiated UGT SNPs, including 4 nonsynonymous SNPs on 4 genes. Uganda exhibits 8.6% of highly divergent UGT SNPs, with 5 significant nonsynonymous SNPs on 4 UGTs (Supplementary Fig. S8 and S9)

We focused our investigation on significantly differentiated non-synonymous SNPs, especially non-synonymous SNPs that could potentially affect the enzyme catalytic activities occurring close to the conserved motifs involved in binding to the glycosyl donor (Fig. 6). Three Nonsynonymous SNPs were detected at the conserved motif of three different UGTs occurring only in susceptible populations (Supplementary Fig. 11) (Table S8). In addition, nonsynonymous changes were detected between sugar donor binding residues 1 and 2 (DBR1 and 2) in *UGT306C2* (c.945T>A, p.His315Gln) only in 3 haplotypes from the resistant Malawi population, and in *UGT302A3* (c.962A>G, p.Asn321Ser) occurring in a frequency of 7 haplotypes from the total 20 haplotypes of the FUMOZ population. Other mutations were detected outside the conserved domain on the signal peptide, transmembrane domain, carboxylic tail, and the N-terminal domain that is believed to determine substrate specificity. Notably, a nonsynonymous SNP on the N-terminal domain of *UGT36C3* (c.226A>G, p.Thr76Ala) that is fixed in the resistant population from Uganda and occurs in 17/20 haplotype of the putatively susceptible populations (Table S8).

**Fig. 6.**
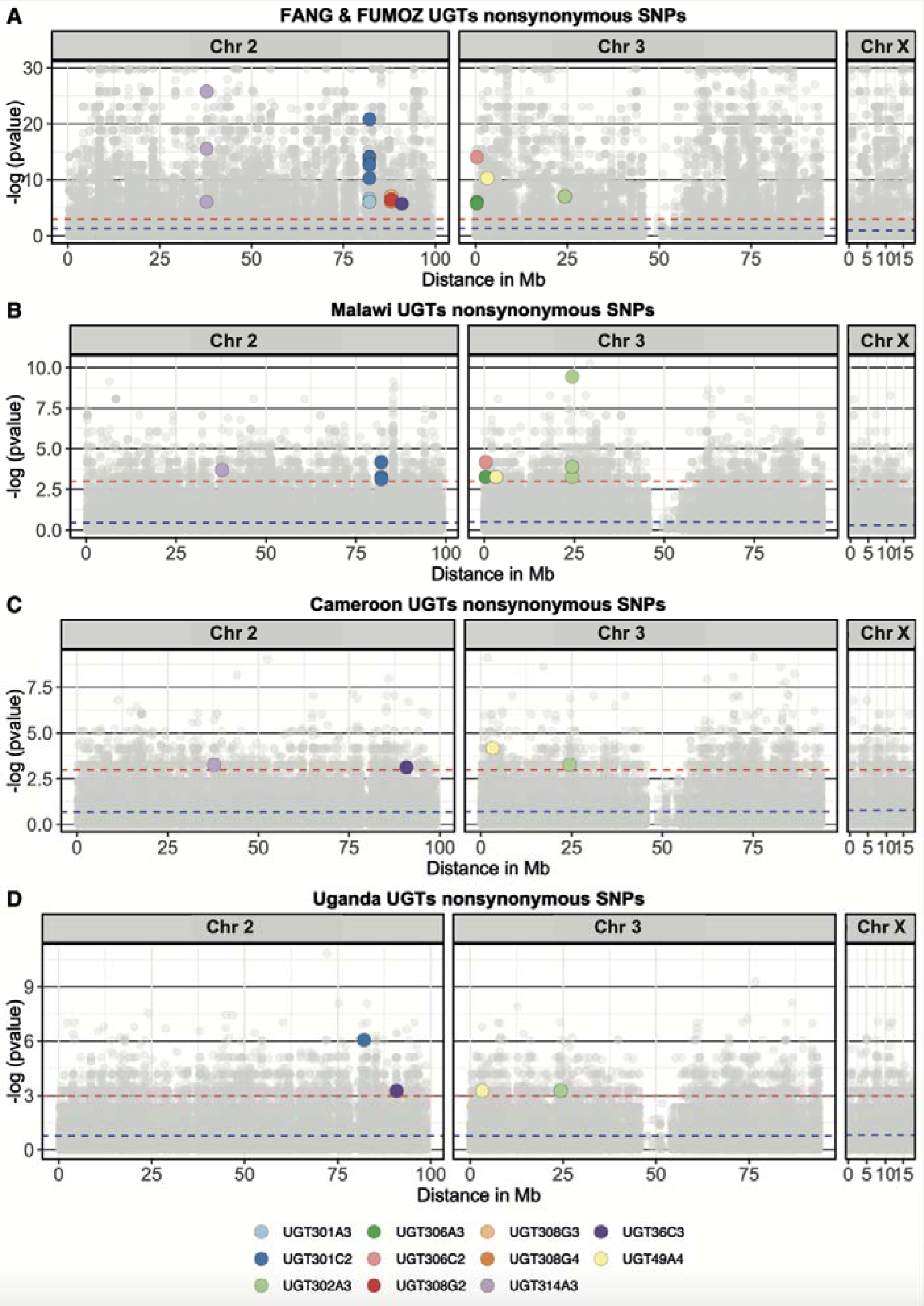
Significantly differentiated UGT genes nonsynonymous SNPs between laboratory colonies and in each country. Non-synonymous SNPs that are significantly differentiated between susceptible and resistant in each analysis are colour-coded according to the associated UGT gene. The red dotted line indicates the significance level (p-value = 0.05). Further details of those nonsynonymous SNPs are included in Table S8 and the location of those SNPs relative to UGTs conserved sites is highlighted in Supplementary Fig. 11.

Overall, the most highly differentiated SNPs in analysis between susceptible in resistant populations from the three countries belong to detoxification genes other than UGTs except for Malawi, where SNPs on *UGT302A3* are the most highly differentiated SNPs (Supplementary Fig. 9) (Supplementary Dataset 7). A nonsynonymous SNP in the N-terminal of *UGT302A3* (c.334C>G, p.Gln112Glu) is amongst the most highly differentiated SNPs in Malawi, occurring in 12 haplotypes of the resistant population and only 1 haplotype of the susceptible population. While in Cameroon the most highly differentiated SNPs in detoxification genes belong to *CYP304B1* and *CYP6AK1* and in Uganda belong to *GSTE7* and *CYP6Z1* (Supplementary Dataset 7).

## 4. Discussion

It is essential to improve our understanding of insecticide resistance since malaria prevention is heavily dependent on insecticide-based interventions. Although other gene families such as cytochrome p450s or GSTs have been extensively studied, the role of UGT genes in pyrethroid resistance remains largely unexplored in malaria vectors. Our study provides a comprehensive genome-wide characterisation of the UGT gene family in *An. funestus* and investigates their expression and selection in geographically distinct populations. A major outcome of the study was the detection of overexpressed and significantly differentiated UGTs in field-collected pyrethroid-resistant populations of *An. funestus*.

### 4.1. Characterisation and evolution of *An. funestus* UGTs

The number of UGT genes identified in *An. funestus* (27) is comparable to that of *An. gambiae* (26), whereas both of *Ae. aegypti* and *D. melanogaster* contain 35 UGT genes. *An. funestus* UGTs exhibit typical characteristics of enzymes that catalyse glycosylation at the C-terminal domain containing significant conservation of 29 amino acids signature motifs and UDP-glycosyl donor binding regions (DBR1 and DBR2), as described for other insects [27, 45]. On the other hand, the N-terminal domain is highly variable between UGTs and is believed to be involved in substrate specificity [38, 44, 46]. The ‘loose’ fit of the substrate binding domain at the N-terminal provides a binding site for structurally diverse substrates by the same UGT isoform [22, 27]. Additionally, a signal peptide is located at the N-terminal indicating that UGTs protein precursor is destined towards the secretory pathway, while the retention of those proteins inside the ER membrane is mediated by the C-terminal hydrophobic transmembrane domain that spans the membrane with a group I topology. Most of the UGT protein resides in the ER and only a small portion of the protein, particularly the cytoplasmic tail, resides in the cytosol [27, 54].

Previous phylogenetic studies of insect UGTs indicate that UGT50 is the only family that is found universally in all insect species [46]. UGT50 family contains one UGT gene from each Diptera species selected for this investigation and the branching pattern of the UGT50 mirrors the species phylogeny among the four Diptera species investigated in this study [46, 76]. The UGT308 family is the largest in *An. funestus* by gene number and in the overall phylogenetic tree and, where 9 *An. funestus* UGTs clustered within this family. The expansion in the UGT308 family was potentially driven by divergence to increase the number of substrates binding for glycosylation and gene duplication, as illustrated by the subfamily UGT308G in *An. funestus* and UGT308J in *Ae. Aegypti*. Based on previous investigations in mosquitoes, genes belonging to the UGT308 family were speculated to be involved in insecticide resistance, and the gene expansion may have evolved to resist consistent exposure to insecticides [38, 77]. However, in our investigation, we did not detect overexpression of *An. funestus* UGT308 genes in resistant field-collected populations.

### 4.2. Overexpression of *UGT310B2* Africa-wide in pyrethroid-resistant populations was detected

In this study, DNA and RNA PoolSeq were analysed to identify an association between *An. funestus* UGTs and pyrethroid resistance. The investigation started by calculating the gene-wise ratio between nonsynonymous and synonymous polymorphisms pN/pS from pooled DNA. The pN/pS genetic test is a powerful population genetic test that requires few assumptions and is a good indicator of selective pressure at a gene level [78]. pN/pS detected what could be a stabilising or a purifying selection acting against changes in the protein sequence in all UGTs from filed collected mosquitoes from Malawi, Cameroon, and Uganda. The pN/pS analysis is limited by only detecting selection pressure in protein-coding regions, however, evolutionary changes in regulatory regions of genes associated with permethrin detoxification may affect the timing and expression level of those genes. Such changes cannot be detected by gene-wise population genetic tests using pooled DNA. Therefore, we investigated the transcriptional regulation of *An. funestus* UGT genes in response to permethrin exposure using total RNA pools. Pooled RNA samples derived from several biologically similar animals, compared to sequencing corresponding individuals individually, reduce the cost of sequencing while retaining similar biological information.

Insecticide resistance surveillance detected an increase in insecticide resistance in Malawi, Cameroon, and Uganda in recent times [7, 9, 13, 17]. Changes in transcriptional regulation in response to permethrin exposure between resistant populations and unexposed populations from the same territory were more evident in Malawi than in Uganda and Cameroon. High resistance levels in Uganda and Cameroon may have obscured the difference in gene expression of detoxification genes induced by permethrin exposure. On the other hand, comparing the transcriptional profile of field-resistant populations and the laboratory-resistant (FUMOZ) to that of the laboratory-susceptible colony (FANG) detected a pronounced difference in detoxification genes expression [17]. We detected overexpression of *UGT310B2* (AFUN011266) across all pyrethroid-resistant populations. UGT overexpression in other mosquito vector populations, such as *An. sinensis* and *A. gambiae*, have been reported and linked to pyrethroid resistance [38, 77]. Although the upregulation of UGTs expression in field-resistant mosquitoes was not as prominent as Phase I detoxification enzymes cytochrome P450s, UGTs still play an important role in the metabolic breakdown of pyrethroid insecticides (Supplementary Fig. S3). UGTs conjugate polar hydrophobic compounds with hydrophilic glycosyl groups, producing more polar metabolites that can be easily excreted from the cell by export transporters [22, 27–29]. Pyrethroids are mostly composed of nonhydrocarbon chains and cyclic easter or acid groups and typically lack a polar group that can be glycosylated by UGTs [79, 80]. Therefore, in the detoxification process of pyrethroids, they undergo initial oxidation by phase I cytochrome P450s, resulting in the production of more polar and reactive metabolites that potentially can be conjugated by UGTs [81].

The overexpression of CYPs compared to other detoxification enzymes indicates their primary role in the detoxification process of pyrethroids. While relative overexpression of UGTs compliments their roles as secondary enzymes interacting with the oxidised pyrethroid, a by-product of CYPs. Previous studies have identified overexpression of *An. funestus* cytochrome P450 genes, including *CYP6P9a*, *CYP6P9b*, *CYP9J11*, *CYP6Z1*, *CYP6M7* and *CYP9K1* in resistant populations. The recombinant protein of those genes expressed *in vitro* metabolises permethrin with significant depletion, therefore reducing the efficacy of permethrin-treated bed nets [7, 8, 14, 17, 25, 26]. The identified overexpression of UGT genes in this study is relevant to current vector control strategies and management. UGTs may contribute to cross-resistance against polar insecticides that can be directly glycosylated enhancing their solubility and excretion. UGTs contribution to resistance against other insecticides that contain a potential glycosylation site has been reported in the Diamondback moth *Plutella xylostella* resistance to chlorantraniliprole [39], the housefly *Musca domestica* resistance to organophosphate [41], and the greenfly cotton aphid *Aphis gossypii* resistance to neonicotinoid and spirotetramat [42, 43].

### 4.3. Targeted sequencing reveals differentiated UGTs and population structure in investigated countries

Sequencing DNA from pools of individuals is a cost-effective approach for conducting population genetics studies, which are otherwise economically challenging to sequence many individual genomes at high coverage [82]. However, to avoid pooling biases and detect SNPs at an individual resolution while maintaining a cost-effective approach, SureSelect target enrichment sequencing was implemented on individual mosquitoes. Polymorphism detected in the targeted region reflects the geographical distance between the collected populations and the FUMOZ reference genome originally from Mozambique (southern Africa). As expected, populations from Southern Africa Malawi and the FANG colony (originally from Angola) harbour lower polymorphism than populations collected from Uganda and Cameroon. The gene-wise differentiation (*F_ST_*) for UGT genes is generally lower than other detoxification genes in all analyses except in Malawi, where *UGT302A3* and *UGT301C2* are the most differentiated detoxification genes, highlighting the potential role of UGTs in detoxification in this region (Supplementary Dataset 5).

A genome-wide process could be inferred from gene-wise Tajima’s D density curves for genes in targeted regions revealing a history of *An. funestus* population in investigated locations, with a likely population expansion after a recent bottleneck effect in Cameroon and Uganda. Gene-wise Tajima’s D density curves for populations from Cameroon and Uganda are skewed towards negative values, whereas populations from southern Africa Malawi, FANG (Angola) and FUMOZ (Mozambique) were close to equilibrium, represented by the coalescent simulation curve. Results in this study, support previous findings of population expansion in the western part of *An. funestus* population range, west of the African Rift Valley [18, 83]. A similar pattern is found in co-distributed malaria vectors *An. gambiae* and *An. coluzzii,* suggesting that those species have responded to common geographic constraints with a population expansion north of the Congo basin and west of the East African Rift Valley [84].

### 4.4. Low-frequency nonsynonymous SNPs within UGTs in field populations show a limited directional selection

Haplotype clustering of UGTs did not reveal a strong directional selection in any of the investigated populations. UGT haplotypes clustered randomly with many singleton haplotypes separated except for UGTs that show significant gene-wise divergence or a potential gene-level selective sweep inferred from Tajima’s D. A dominant haplotype was detected in *UGT301C2* that is significantly differentiated gene-wise in analysis between resistant and susceptible populations in Malawi and Africa-wide. A potential gene-level selective sweep is detected in Malawi at *UGT301C2*, while in Cameroon the significantly negative Tajima’s D is not a deviation from the genome-wide trend. Haplotype clustering was detected for *UGT306A3* and C2 both genes recorded significant Tajima’s D in Cameroon while *UGT306C2* is significantly differentiated between resistant and putatively susceptible populations from Uganda. The haplotype clustering for *UGT301C2* and *UGT306A3* revealed limited directional selection with a predominant haplotype detected in field-collected populations, while *UGT306C2* haplotypes from filed populations cluster in more than one node. The limited selection in UGTs might reflect their role in pyrethroid detoxification as phase II enzymes in the detoxification pathway of pyrethroid compared to the previously described selection for *CYP9K1* in Uganda, and *CYP6P9A* and *CYP6P9B* in populations from southern Africa (Supplementary Fig. 10) [7, 10, 11, 14].

The resulting averaged values of gene-wise *F_ST_* can mask or lower the magnitude of a positive selection where a combination of positive and negative selection within the gene at different times along its evolution may cancel each other out. Most of the significantly differentiated SNPs in all investigated populations are synonymous and intronic. Therefore, we focused the investigation on significantly differentiated nonsynonymous at the conserved motifs. Nonsynonymous mutations causing amino acid changes at conserved motifs acting as a binding site to the UDP-glycosyl donor are found in low frequencies in field-collected populations but with a significant differentiation between putatively susceptible and resistant populations. Other nonsynonymous mutations were detected outside the conserved motifs at the signal peptide, transmembrane domain, carboxylic tail, and the N-terminal domain.

Changes in the N-terminal domain might have a detrimental effect on substrate specificity. However, the N-terminal domain generally exhibits greater sequence divergence between UGT isoforms to enable the glycosylation of diverse substrates by the same UGT isoform [22, 27]. Research on allelic variations in human UGTs indicated that amino acid substitutions at key residues can affect the enzyme-substrate selectivity, glycosylation activity and overall drug metabolism [85–88]. Unless those nonsynonymous mutations identified in this research are validated using recombinant proteins substantiating their functional role in pyrethroid detoxification is challenging. To capture the complete effect of those nonsynonymous sites in UGTs activity we recommend using a baculovirus expression system in insect cells to account for post-translational modifications that have a significant impact on their activity [89]. Post-translational modification of UGTs specifically *N*-glycosylation was demonstrated to be important for UGT protein folding and catalytic activity [85, 86, 89]. Post-translational modification of UGTs specifically *N*-glycosylation was demonstrated to be important for UGT protein folding and catalytic activity [85, 86].

It has been established that direct selection of nonsynonymous SNPs causing amino acid alterations in *An. funestus* detoxification genes such as P450s [7, 8, 90] and GSTs [15] enhance their detoxification activities. Despite the low frequency of the UGTs nonsynonymous SNPs outlined in this paper they are the first nonsynonymous SNPs to be reported in *An. funestus* UGTs. The selection detected in *An. funestus* UGTs nonsynonymous SNPs is limited and further illustrate the secondary role of *An. funestus* UGTs in pyrethroid resistance. However, the outlined significantly differentiated UGTs nonsynonymous SNPs will help future prediction of cross-resistance to insecticides that can be directly detoxicated by UGTs.

## Conclusion

In this study, we have highlighted the potential role of UGTs in pyrethroid resistance in resistant laboratory colonies and field-collected *An. funestus* populations. Notably, we have identified a common overexpression of *UGT310B2* across various regions in Africa. The overexpression of *UGT310B2* in the FUMOZ colony was confirmed using quantitative PCR. The gene-wise (*F_ST_*) differentiation for UGT genes is generally lower than other detoxification genes in all analyses except in Malawi, where *UGT302A3* and *UGT301C2* are the most differentiated detoxification genes, highlighting the potential role of UGTs in detoxification in this region. In addition, SNPs belonging to *UGT302A3* were the most highly differentiated SNPs in Malawi between resistant and putatively susceptible based on p-values quantifying differences in allele frequencies (*pF_ST_*). In Uganda, a significant gene-wise *F_ST_* differentiation was detected in *UGT306C2*. The high gene-wise *F_ST_* divergence of *UGT301C2* and *UGT306C2* was supported by a limited selection with limited clustering of haplotypes from those regions in the haplotype tree. Gene-wise Tajima’s D density curves infer genome-wide process and reveal population structures of *An. funestus* population from the three countries, supporting previous observations. In addition, this study identified significantly differentiated UGTs nonsynonymous SNPs that might implicate UGTs detoxification activities and produced detailed records of those SNPs. Findings in this study have important implications for current vector control strategies and management as UGT enzymes may confer cross-resistance to other polar insecticides, which can be directly detoxified by UGTs. Further investigations using more recent field-collected populations across Africa should be carried out to explore the role of UGT genes in pyrethroid resistance and predict their potential for cross-resistance to other insecticides.

## Data availability

Read data of pooled whole genome sequencing PoolSeq are available in European Nucleotide Archive under accessions (PRJEB24379, PRJEB13485 and PRJEB24384). SureSelect data are available under accession numbers PRJEB24520 (Cameroon), PRJEB47287 (Malawi and Uganda) and PRJEB24506 (FANG colony). RNA-seq data are available under accession number PRJEB24351 and PRJEB10294.

## Authorship contribution statement

**Talal Al-Yazeedi:** The design of the research, the experimental work, data analysis, data curation, data visualisation and the preparation of the manuscript. **Abdullahi Muhammad:** Experimental work, reviewing and editing of the manuscript. **Seung-Joon Ahn:** Data analysis, reviewing and editing of the manuscript**. Helen Irving:** Data collection. **Jack Hearn:** The design of the research, data analysis, data collection and reviewing and editing of the manuscript. **Charles S. Wondji:** The design of the research, funding acquisition and reviewing and editing of the manuscript.

## Acknowledgments

This work was supported by a Wellcome Trust Senior Research Fellowships in Biomedical Sciences to Charles S. Wondji (101893/Z/13/Z and 217188/Z/19/Z) and a Bill and Melinda Gates Foundation grant to CSW (INV-006003). For the purpose of open access, the authors have applied a CC BY public copyright licence to any Author Accepted Manuscript version arising from this submission.

The funders had no role in study design, data collection and analysis, decision to publish, or preparation of the manuscript.

## References

1. Organization, W.H., World malaria report 2022. 2022: World Health Organization.

2. Bhatt, S., et al., The effect of malaria control on Plasmodium falciparum in Africa between 2000 and 2015. Nature, 2015. 526(7572): p. 207–211.

3. Hemingway, J., The way forward for vector control. Science, 2017. 358(6366): p. 998–999.

4. Hemingway, J., et al., Averting a malaria disaster: will insecticide resistance derail malaria control? Lancet, 2016. 387(10029): p. 1785–8.

5. Sinka, M.E., et al., The dominant Anopheles vectors of human malaria in Africa, Europe and the Middle East: occurrence data, distribution maps and bionomic précis. Parasit Vectors, 2010. 3: p. 117.

6. Atoyebi, S.M., et al., Investigating the molecular basis of multiple insecticide resistance in a major malaria vector Anopheles funestus (sensu stricto) from Akaka-Remo, Ogun State, Nigeria. Parasites & Vectors, 2020. 13(1): p. 423.

7. Hearn, J., et al., Multi-omics analysis identifies a CYP9K1 haplotype conferring pyrethroid resistance in the malaria vector Anopheles funestus in East Africa. Mol Ecol, 2022. 31(13): p. 3642–3657.

8. Ibrahim, S.S., et al., The P450 CYP6Z1 confers carbamate/pyrethroid cross-resistance in a major African malaria vector beside a novel carbamate-insensitive N485I acetylcholinesterase-1 mutation. Mol Ecol, 2016. 25(14): p. 3436–52.

9. Morgan, J.C., et al., Pyrethroid resistance in an Anopheles funestus population from Uganda. PLoS One, 2010. 5(7): p. e11872.

10. Mugenzi, L.M.J., et al., A 6.5-kb intergenic structural variation enhances P450-mediated resistance to pyrethroids in malaria vectors lowering bed net efficacy. Molecular Ecology, 2020. 29(22): p. 4395–4411.

11. Mugenzi, L.M.J., et al., Cis-regulatory CYP6P9b P450 variants associated with loss of insecticide-treated bed net efficacy against Anopheles funestus. Nat Commun, 2019. 10(1): p. 4652.

12. Mulamba, C., et al., Widespread Pyrethroid and DDT Resistance in the Major Malaria Vector Anopheles funestus in East Africa Is Driven by Metabolic Resistance Mechanisms. PLOS ONE, 2014. 9(10): p. e110058.

13. Riveron, J.M., et al., Rise of multiple insecticide resistance in Anopheles funestus in Malawi: a major concern for malaria vector control. Malar J, 2015. 14: p. 344.

14. Riveron, J.M., et al., Directionally selected cytochrome P450 alleles are driving the spread of pyrethroid resistance in the major malaria vector Anopheles funestus. Proceedings of the National Academy of Sciences, 2013. 110(1): p. 252–257.

15. Riveron, J.M., et al., A single mutation in the GSTe2 gene allows tracking of metabolically based insecticide resistance in a major malaria vector. Genome biology, 2014. 15(2): p. R27–R27.

16. Tchigossou, G., et al., Molecular basis of permethrin and DDT resistance in an Anopheles funestus population from Benin. Parasites & Vectors, 2018. 11(1): p. 602.

17. Weedall, G.D., et al., A cytochrome P450 allele confers pyrethroid resistance on a major African malaria vector, reducing insecticide-treated bednet efficacy. Sci Transl Med, 2019. 11(484).

18. Weedall, G.D., et al., An Africa-wide genomic evolution of insecticide resistance in the malaria vector Anopheles funestus involves selective sweeps, copy number variations, gene conversion and transposons. PLOS Genetics, 2020. 16(6): p. e1008822.

19. Wondji, C.S., et al., Two duplicated P450 genes are associated with pyrethroid resistance in Anopheles funestus, a major malaria vector. Genome Res, 2009. 19(3): p. 452–9.

20. Jacob, M.R., et al., Insecticide Resistance in Malaria Vectors: An Update at a Global Scale, in Towards Malaria Elimination, M. Sylvie and D. Vas, Editors. 2018, IntechOpen: Rijeka. p. Ch. 7.

21. Hemingway, J., et al., The molecular basis of insecticide resistance in mosquitoes. Insect Biochem Mol Biol, 2004. 34(7): p. 653–65.

22. Bock, K.W., Vertebrate UDP-glucuronosyltransferases: functional and evolutionary aspects. Biochemical Pharmacology, 2003. 66(5): p. 691–696.

23. Black, W.C.t., et al., From Global to Local-New Insights into Features of Pyrethroid Detoxification in Vector Mosquitoes. Insects, 2021. 12(4).

24. Grant, D.M., Detoxification pathways in the liver. J Inherit Metab Dis, 1991. 14(4): p. 421–30.

25. Riveron, J.M., et al., Genome-Wide Transcription and Functional Analyses Reveal Heterogeneous Molecular Mechanisms Driving Pyrethroids Resistance in the Major Malaria Vector Anopheles funestus Across Africa. G3 Genes|Genomes|Genetics, 2017. 7(6): p. 1819–1832.

26. Wondji, C.S., et al., RNAseq-based gene expression profiling of the Anopheles funestus pyrethroid-resistant strain FUMOZ highlights the predominant role of the duplicated CYP6P9a/b cytochrome P450s. G3 Genes|Genomes|Genetics, 2021. 12(1).

27. Meech, R. and P.I. Mackenzie, Structure and function of uridine diphosphate glucuronosyltransferases. Clin Exp Pharmacol Physiol, 1997. 24(12): p. 907–15.

28. Burchell, B. and M.W.H. Coughtrie, UDP-glucuronosyltransferases. Pharmacology & Therapeutics, 1989. 43(2): p. 261–289.

29. Meech, R., et al., The UDP-Glycosyltransferase (UGT) Superfamily: New Members, New Functions, and Novel Paradigms. Physiol Rev, 2019. 99(2): p. 1153–1222.

30. Després, L., J.P. David, and C. Gallet, The evolutionary ecology of insect resistance to plant chemicals. Trends Ecol Evol, 2007. 22(6): p. 298–307.

31. Wiesen, B., et al., Sequestration of host-plant-derived flavonoids by lycaenid butterflyPolyommatus icarus. J Chem Ecol, 1994. 20(10): p. 2523–38.

32. T L Hopkins, a. and K.J. Kramer, Insect Cuticle Sclerotization. Annual Review of Entomology, 1992. 37(1): p. 273–302.

33. Luque, T., K. Okano, and D.R. O’Reilly, Characterization of a novel silkworm (Bombyx mori) phenol UDP-glucosyltransferase. European Journal of Biochemistry, 2002. 269(3): p. 819–825.

34. Ahmad, S.A. and T.L. Hopkins, β-Glucosylation of plant phenolics by phenol β-glucosyltransferase in larval tissues of the tobacco hornworm, Manduca sexta (L.). Insect Biochemistry and Molecular Biology, 1993. 23(5): p. 581–589.

35. Kojima, W., et al., Physiological adaptation of the Asian corn borer Ostrinia furnacalis to chemical defenses of its host plant, maize. Journal of Insect Physiology, 2010. 56(9): p. 1349–1355.

36. Cui, X., et al., Molecular Mechanism of the UDP-Glucuronosyltransferase 2B20-like Gene (AccUGT2B20-like) in Pesticide Resistance of Apis cerana cerana. Frontiers in Genetics, 2020. 11.

37. Hu, B., et al., The expression of Spodoptera exigua P450 and UGT genes: tissue specificity and response to insecticides. Insect Science, 2019. 26(2): p. 199–216.

38. Zhou, Y., et al., UDP-glycosyltransferase genes and their association and mutations associated with pyrethroid resistance in Anopheles sinensis (Diptera: Culicidae). Malar J, 2019. 18(1): p. 62.

39. Li, X., et al., Over-expression of UDP–glycosyltransferase gene UGT2B17 is involved in chlorantraniliprole resistance in Plutella xylostella (L.). Pest Management Science, 2017. 73(7): p. 1402–1409.

40. Grant, C., et al., Overexpression of the UDP-glycosyltransferase UGT34A23 confers resistance to the diamide insecticide chlorantraniliprole in the tomato leafminer, Tuta absoluta. Insect Biochemistry and Molecular Biology, 2023. 159: p. 103983.

41. Lee, S.-W., et al., Metabolic resistance mechanisms of the housefly (Musca domestica) resistant to pyraclofos. Pesticide Biochemistry and Physiology, 2006. 85(2): p. 76–83.

42. Chen, X., et al., UDP-glucosyltransferases potentially contribute to imidacloprid resistance in Aphis gossypii glover based on transcriptomic and proteomic analyses. Pesticide Biochemistry and Physiology, 2019. 159: p. 98–106.

43. Pan, Y., et al., UDP-glycosyltransferases contribute to spirotetramat resistance in Aphis gossypii Glover. Pestic Biochem Physiol, 2020. 166: p. 104565.

44. Li, X., et al., Characterization of UDP-glucuronosyltransferase genes and their possible roles in multi-insecticide resistance in Plutella xylostella (L.). Pest Management Science, 2018. 74(3): p. 695–704.

45. Ahn, S.-J., H. Vogel, and D.G. Heckel, Comparative analysis of the UDP-glycosyltransferase multigene family in insects. Insect Biochemistry and Molecular Biology, 2012. 42(2): p. 133–147.

46. Huang, F.-F., et al., The UDP-glucosyltransferase multigene family in Bombyx mori. BMC Genomics, 2008. 9(1): p. 563.

47. Ahn, S.-J. and S.J. Marygold, The UDP-Glycosyltransferase Family in Drosophila melanogaster: Nomenclature Update, Gene Expression and Phylogenetic Analysis. Frontiers in Physiology, 2021. 12.

48. Mistry, J., et al., Pfam: The protein families database in 2021. Nucleic Acids Research, 2021. 49(D1): p. D412–D419.

49. Blum, M., et al., The InterPro protein families and domains database: 20 years on. Nucleic Acids Research, 2021. 49(D1): p. D344–D354.

50. Giraldo-Calderón, G.I., et al., VectorBase: an updated bioinformatics resource for invertebrate vectors and other organisms related with human diseases. Nucleic Acids Research, 2015. 43(D1): p. D707–D713.

51. Ghurye, J., et al., A chromosome-scale assembly of the major African malaria vector Anopheles funestus. Gigascience, 2019. 8(6).

52. Krogh, A., et al., Predicting transmembrane protein topology with a hidden Markov model: application to complete genomes. J Mol Biol, 2001. 305(3): p. 567–80.

53. Price, M.N., P.S. Dehal, and A.P. Arkin, FastTree: Computing Large Minimum Evolution Trees with Profiles instead of a Distance Matrix. Molecular Biology and Evolution, 2009. 26(7): p. 1641–1650.

54. Mackenzie, P.I., et al., The UDP glycosyltransferase gene superfamily: recommended nomenclature update based on evolutionary divergence. Pharmacogenetics, 1997. 7(4): p. 255–69.

55. Danecek, P., et al., Twelve years of SAMtools and BCFtools. GigaScience, 2021. 10(2).

56. Edgar, R.C., MUSCLE: multiple sequence alignment with high accuracy and high throughput. Nucleic Acids Res, 2004. 32(5): p. 1792–7.

57. Leigh, J.W. and D. Bryant, popart: full-feature software for haplotype network construction. Methods in Ecology and Evolution, 2015. 6(9): p. 1110–1116.

58. Hearn, J., et al., Gene Conversion Explains Elevated Diversity in the Immunity Modulating APL1 Gene of the Malaria Vector Anopheles funestus. Genes (Basel), 2022. 13(6).

59. Hunt, R.H., et al., Laboratory selection for and characteristics of pyrethroid resistance in the malaria vector Anopheles funestus. Med Vet Entomol, 2005. 19(3): p. 271–5.

60. Gillies, M.T. and M. Coetzee, A supplement to the Anophelinae of Africa South of the Sahara. Publ S Afr Inst Med Res, 1987. 55: p. 1–143.

61. Koekemoer, L.L., et al., A cocktail polymerase chain reaction assay to identify members of the Anopheles funestus (Diptera: Culicidae) group. Am J Trop Med Hyg, 2002. 66(6): p. 804–11.

62. Wondji, C.S., et al., Mapping a Quantitative Trait Locus (QTL) conferring pyrethroid resistance in the African malaria vector Anopheles funestus. BMC Genomics, 2007. 8(1): p. 34.

63. Robinson, M.D., D.J. McCarthy, and G.K. Smyth, edgeR: a Bioconductor package for differential expression analysis of digital gene expression data. Bioinformatics, 2010. 26(1): p. 139–40.

64. Schmittgen, T.D. and K.J. Livak, Analyzing real-time PCR data by the comparative CT method. Nature Protocols, 2008. 3(6): p. 1101–1108.

65. Nelson, C.W., L.H. Moncla, and A.L. Hughes, SNPGenie: estimating evolutionary parameters to detect natural selection using pooled next-generation sequencing data. Bioinformatics, 2015. 31(22): p. 3709–11.

66. Li, H. and R. Durbin, Fast and accurate short read alignment with Burrows-Wheeler transform. Bioinformatics, 2009. 25(14): p. 1754–60.

67. Li, H., et al., The Sequence Alignment/Map format and SAMtools. Bioinformatics (Oxford, England), 2009. 25(16): p. 2078–2079.

68. McKenna, A., et al., The Genome Analysis Toolkit: a MapReduce framework for analyzing next-generation DNA sequencing data. Genome research, 2010. 20(9): p. 1297–1303.

69. Garrison E, M.G., Haplotype-based variant detection from short-read sequencing. arXiv preprint arXiv, 2012. 1207.3907(q-bio.GN).

70. Garrison, E., et al., Vcflib and tools for processing the VCF variant call format. bioRxiv, 2021: p. 2021.05.21.445151.

71. Cingolani, P., et al., A program for annotating and predicting the effects of single nucleotide polymorphisms, SnpEff: SNPs in the genome of Drosophila melanogaster strain w1118; iso-2; iso-3. Fly (Austin), 2012. 6(2): p. 80–92.

72. Pfeifer, B., et al., PopGenome: an efficient Swiss army knife for population genomic analyses in R. Molecular biology and evolution, 2014. 31(7): p. 1929–1936.

73. Nielsen, R., et al., Genomic scans for selective sweeps using SNP data. Genome Res, 2005. 15(11): p. 1566–75.

74. Hudson, R.R., Generating samples under a Wright–Fisher neutral model of genetic variation. Bioinformatics, 2002. 18(2): p. 337–338.

75. Ayala, D., et al., The genome sequence of the malaria mosquito, Anopheles funestus, Giles, 1900 [version 1; peer review: 2 approved]. Wellcome Open Research, 2022. 7(287).

76. Neafsey, D.E., et al., Mosquito genomics. Highly evolvable malaria vectors: the genomes of 16 Anopheles mosquitoes. Science, 2015. 347(6217): p. 1258522.

77. Nkya, T.E., et al., Insecticide resistance mechanisms associated with different environments in the malaria vector Anopheles gambiae: a case study in Tanzania. Malar J, 2014. 13: p. 28.

78. Yang, Z. and J.P. Bielawski, Statistical methods for detecting molecular adaptation. Trends Ecol Evol, 2000. 15(12): p. 496–503.

79. Khambay, B.P.S. and P.J. Jewess, 6.1 - Pyrethroids, in Comprehensive Molecular Insect Science, L.I. Gilbert, Editor. 2005, Elsevier: Amsterdam. p. 1–29.

80. Soderlund, D.M., Molecular mechanisms of pyrethroid insecticide neurotoxicity: recent advances. Arch Toxicol, 2012. 86(2): p. 165–81.

81. Guengerich, F.P., Cytochrome P450 research and The Journal of Biological Chemistry. J Biol Chem, 2019. 294(5): p. 1671–1680.

82. Cutler, D.J. and J.D. Jensen, To pool, or not to pool? Genetics, 2010. 186(1): p. 41–3.

83. Michel, A.P., et al., Rangewide population genetic structure of the African malaria vector Anopheles funestus. Molecular Ecology, 2005. 14(14): p. 4235–4248.

84. Genetic diversity of the African malaria vector Anopheles gambiae. Nature, 2017. 552(7683): p. 96–100.

85. Nakajima, M., et al., N-Glycosylation plays a role in protein folding of human UGT1A9. Biochemical Pharmacology, 2010. 79(8): p. 1165–1172.

86. Nakamura, T., et al., Introduction of an N-Glycosylation Site into UDP-Glucuronosyltransferase 2B3 Alters Its Sensitivity to Cytochrome P450 3A1-Dependent Modulation. Front Pharmacol, 2016. 7: p. 427.

87. Korprasertthaworn, P., et al., Effects of amino acid substitutions at positions 33 and 37 on UDP-glucuronosyltransferase 1A9 (UGT1A9) activity and substrate selectivity. Biochem Pharmacol, 2012. 84(11): p. 1511–21.

88. Kim, J.Y., et al., Comprehensive variant screening of the UGT gene family. Yonsei Med J, 2014. 55(1): p. 232–9.

89. Chambers, A.C., et al., Overview of the Baculovirus Expression System. Curr Protoc Protein Sci, 2018. 91: p. 5.4.1–5.4.6.

90. Ibrahim, S.S., et al., Allelic Variation of Cytochrome P450s Drives Resistance to Bednet Insecticides in a Major Malaria Vector. PLOS Genetics, 2015. 11(10): p. e1005618.1.

