## Supplementary Figures for "Overexpression and nonsynonymous mutations of UDP-glycosyltransferases potentially associated with pyrethroid resistance in *Anopheles funestus*"

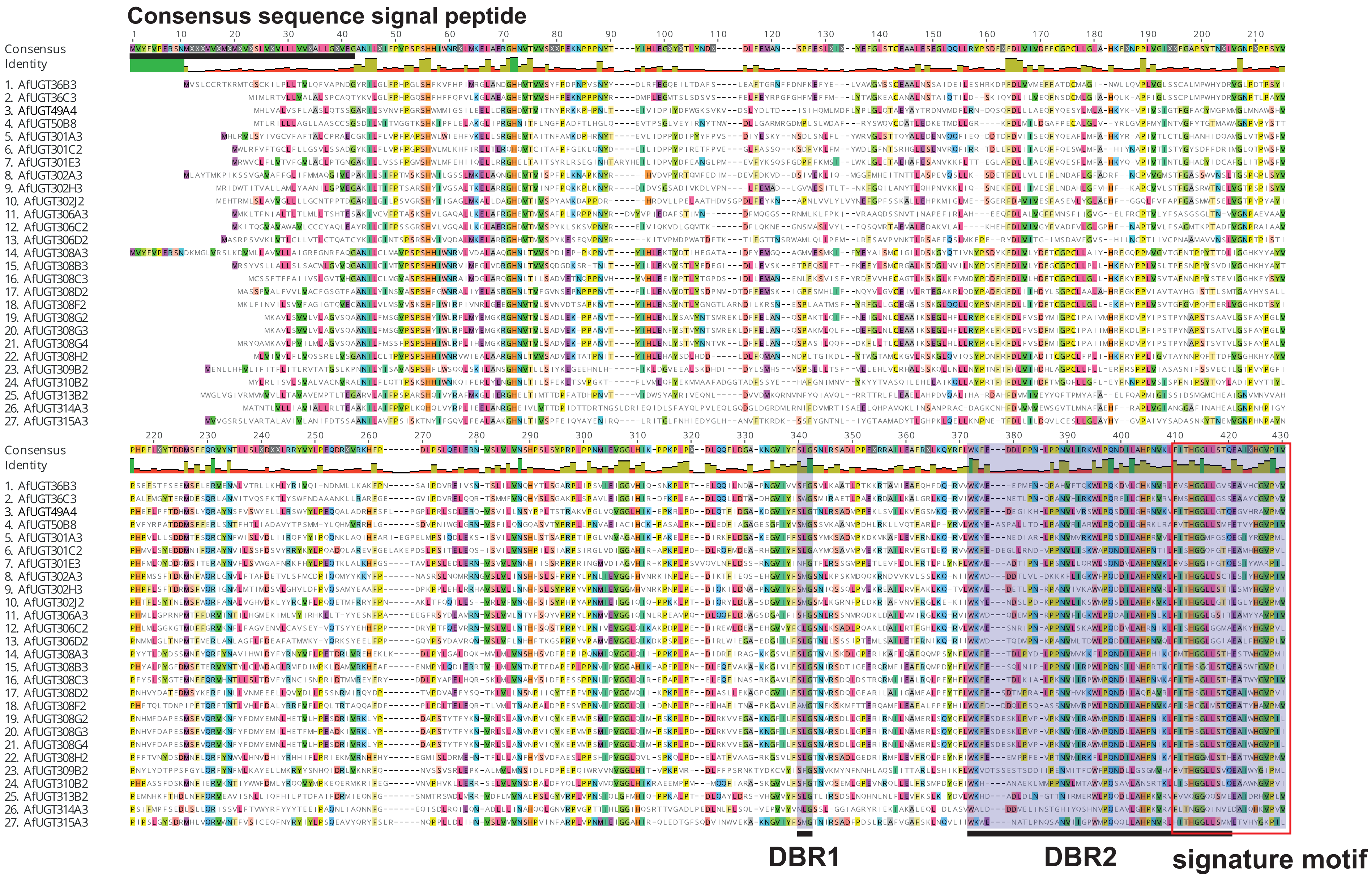
Consensus sequence signal peptide
Consensus
Identity
1. AfUGT36B3
2. AfUGT36C3
3. AfUGT49A4
4. AfUGT50B8
5. AfUGT301A3
6. AfUGT301C2
7. AfUGT301E3
8. AfUGT302A3
9. AfUGT302H3
10. AfUGT302J2
11. AfUGT306A3
12. AfUGT306C2
13. AfUGT306D2
14. AfUGT308A3
15. AfUGT308B3
16. AfUGT308C3
17. AfUGT308D2
18. AfUGT308F2
19. AfUGT308G2
20. AfUGT308G3
21. AfUGT308G4
22. AfUGT308H2
23. AfUGT309B2
24. AfUGT310B2
25. AfUGT313B2
26. AfUGT314A3
27. AfUGT315A3
DBR1
DBR2
signature motif

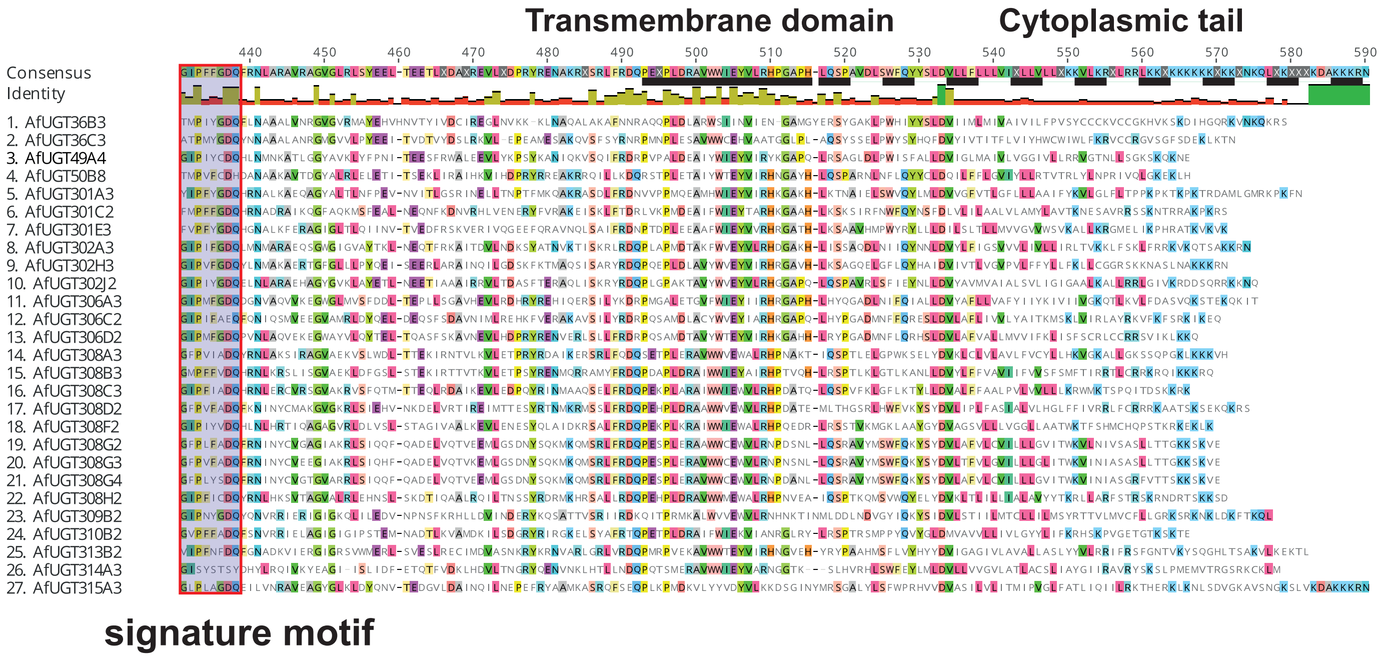

**Fig. S1. Alignment of *An. funestus* UGTs amino acids**. Alignment reveals that the C-terminal domain is more conserved than the highly variable N-terminal domain. The 29 amino acids sequence long signature motif on the C-terminal domain is highlighted by a red box. Conserved signature motifs including the DBR1 and the DBR2 motifs are highlighted in purple.

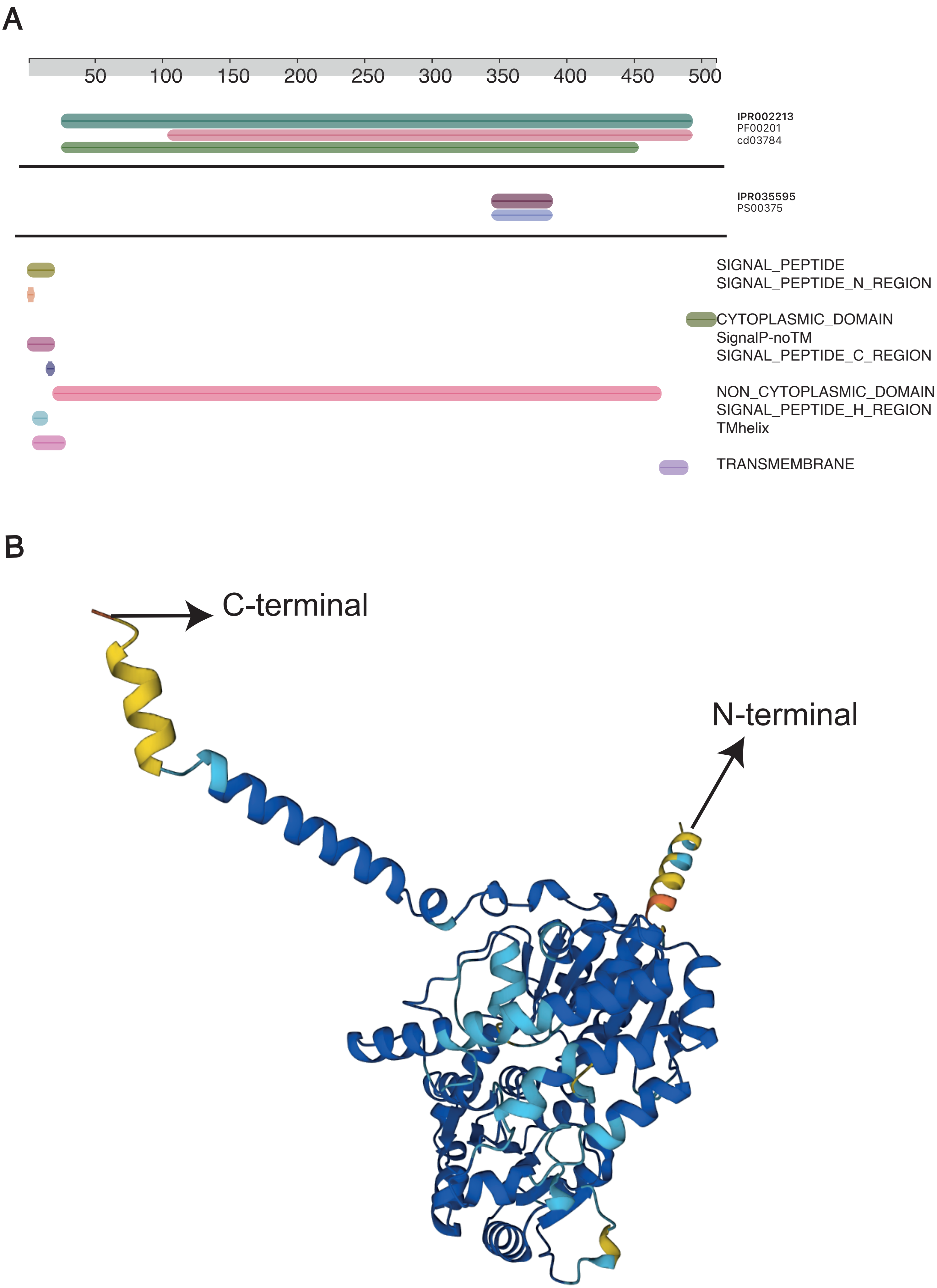

A
50
100
150
200
250
300
350
400
450
500
IPR002213
PF00201
cd03784
IPR035595
PS00375
SIGNAL_PEPTIDE
SIGNAL_PEPTIDE_N_REGION
CYTOPLASMIC_DOMAIN
SignalP-noTM
SIGNAL_PEPTIDE_C_REGION
NON_CYTOPLASMIC_DOMAIN
SIGNAL_PEPTIDE_H_REGION
TMhelix
TRANSMEMBRANE
B
C-terminal
N-terminal

**Fig. S2. Protein function analysis of UGT50B8.** *An funestus* UGT50B8 was selected for functional analysis because UGT50 is the only family that is found universally in all insect species. Protein functional analysis conducted by InterPro scan http://www.ebi.ac.uk/interpro/search/sequence/ indicates that the protein pfam Id is PF00201 belonging to the UDP glycosyltransferases superfamily **(A)**. The functional analysis locates the UGTs conserved motif in the C- terminal domain. The majority of the protein is non-cytoplasmic resides in the endoplasmic reticulum (ER) while a small portion of the protein is cytoplasmic. The retention of the protein inside the ER is mediated by hydrophobic trans-membrane domain **(A)**. The protein structure was predicted using AlphaFold **(B)**. AlphaFold structures are colored using a per-residue confidence metric called pLDDT, which is scaled from 1 - 100. **Very high** (pLDDT > 90) **Confident** (90 > pLDDT > 70) **Low** (70 > pLDDT > 50) **Very low** (pLDDT < 50)

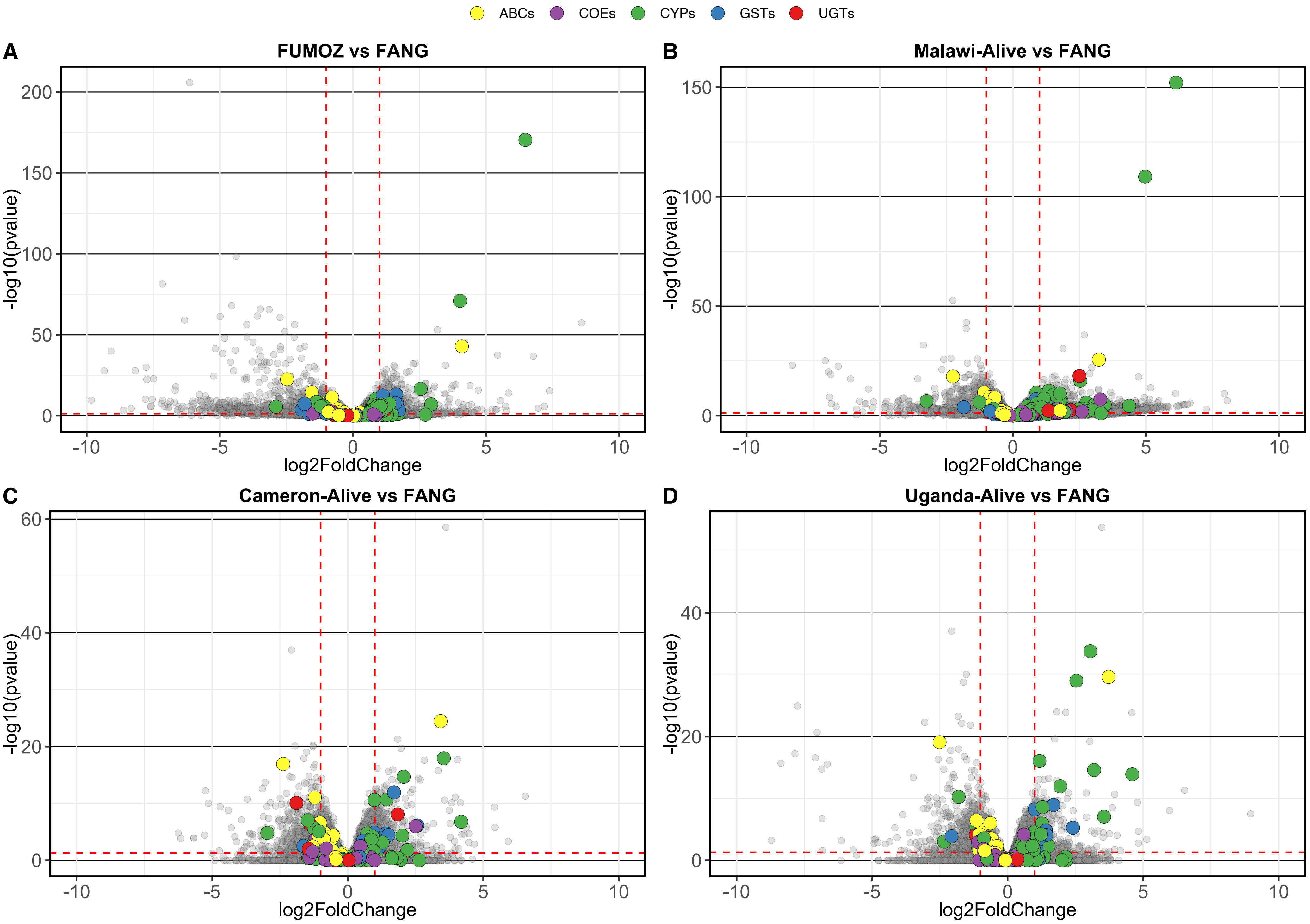

ABCs
COEs
CYPs
GSTs
UGTs
A
FUMOZ vs FANG
B
Malawi-Alive vs FANG
C
Cameron-Alive vs FANG
D
Uganda-Alive vs FANG
-log10(pvalue)
log2FoldChange

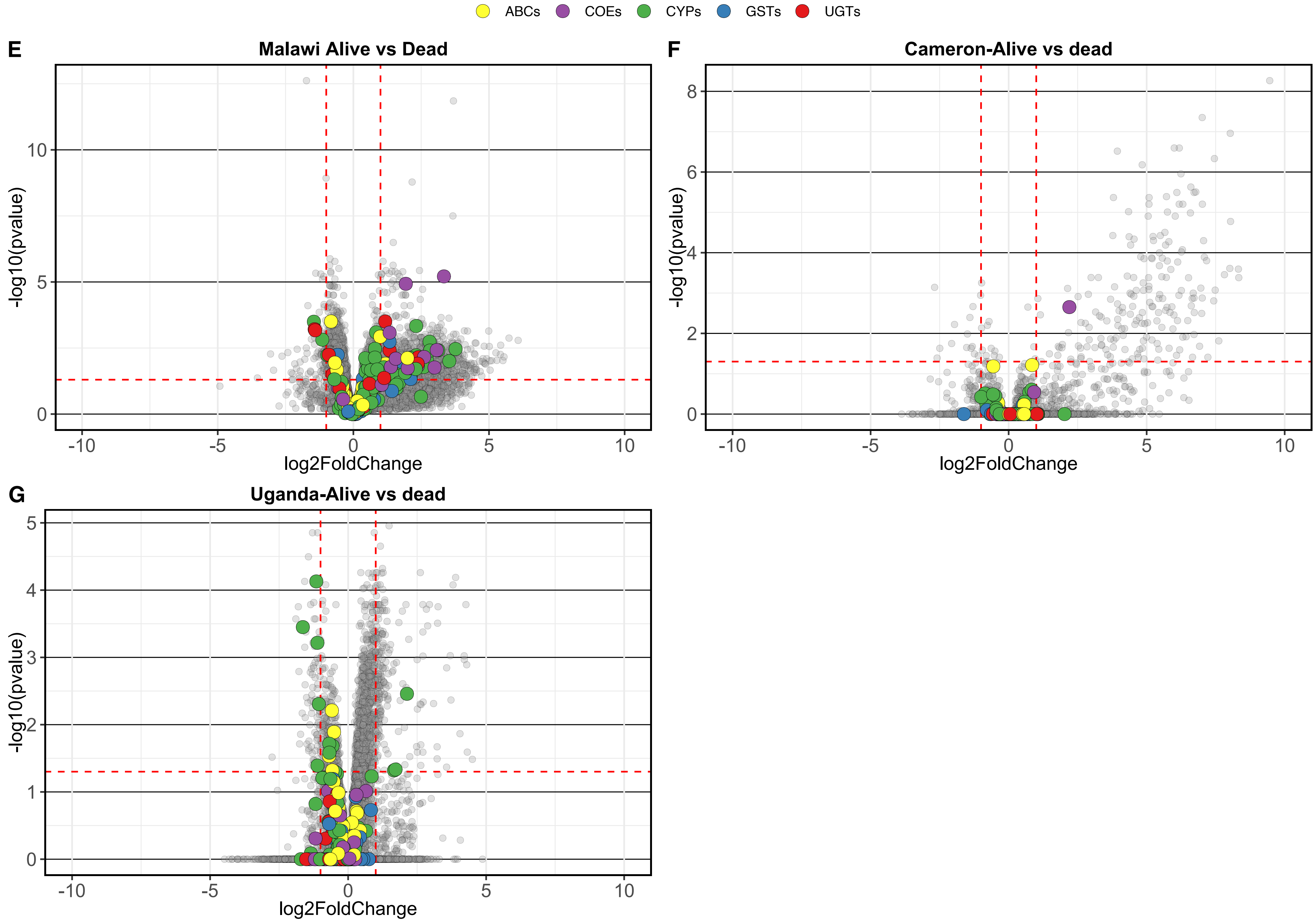
ABCs
COEs
CYPs
GSTs
UGTs
E
Malawi Alive vs Dead
-log10(pvalue)
log2FoldChange
F
Cameron-Alive vs dead
-log10(pvalue)
log2FoldChange
G
Uganda-Alive vs dead
-log10(pvalue)
log2FoldChange

**Fig. S3. Detoxification genes differential expression analysis in Malawi, Uganda, Cameroon and the resistant colony FUMOZ.** The transcription profiles of the resistant colony FUMOZ, filed-collected resistant mosquitoes from Malawi, Uganda and Cameroon was compared to the FANG transcription profile **(A-D)**. In addition transactional profiles of filed-collected resistant population was compared to the transactional profile of a population of mosquitoes unexposed to permeation from the same  same country **(E-G)**. Differentially regulated genes were defined as those with a corrected p-value threshold of < 0.05 and log2(fold change) > 1 indicated by the dotted red line in each plot. Differentially expressed detoxification genes were color coded according to the plot legend.

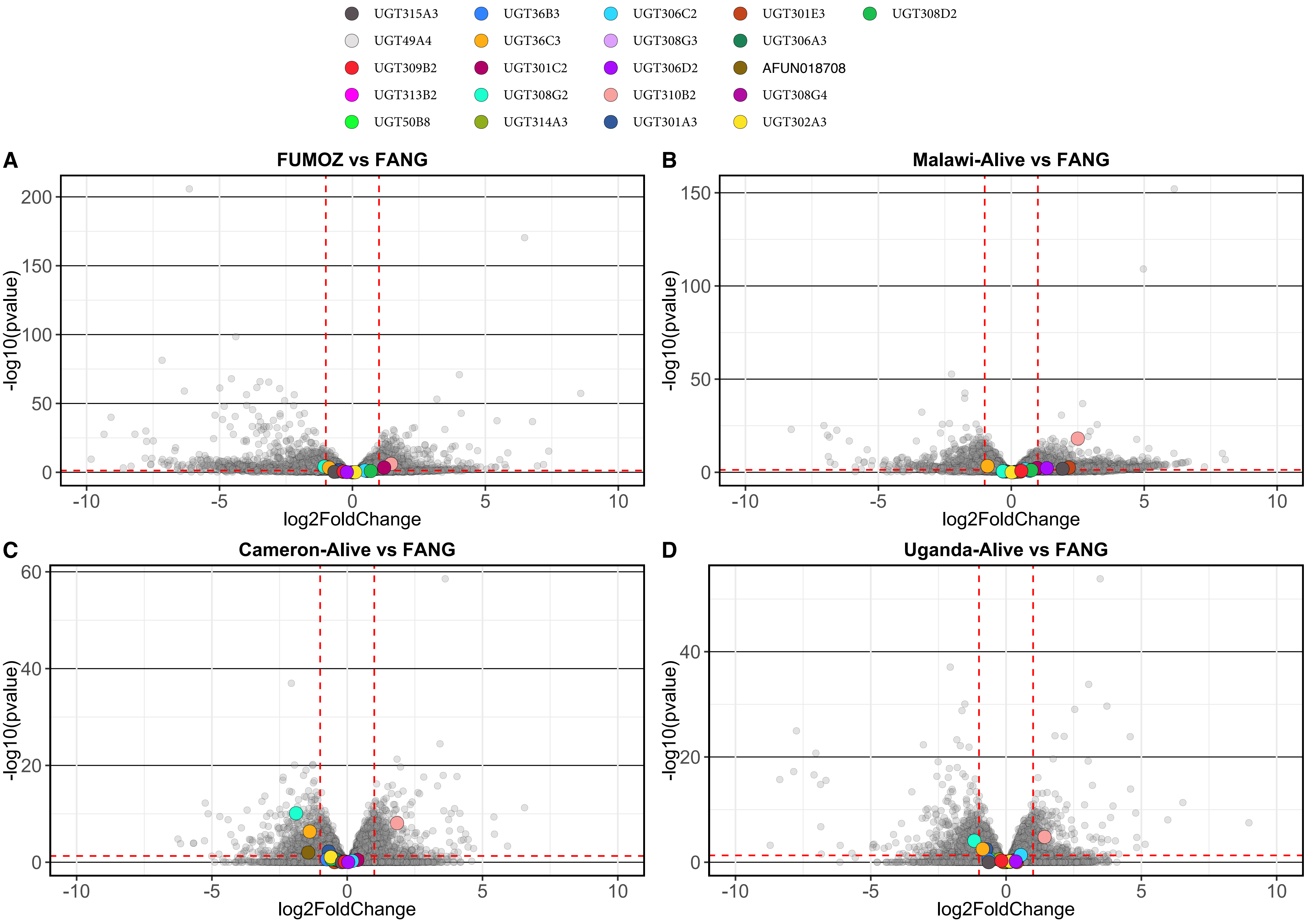

UGT315A3
UGT49A4
UGT309B2
UGT313B2
UGT50B8
UGT36B3
UGT36C3
UGT301C2
UGT308G2
UGT314A3
UGT306C2
UGT308G3
UGT306D2
UGT310B2
UGT301A3
UGT301E3
UGT306A3
AFUN018708
UGT308G4
UGT302A3
UGT308D2
A
FUMOZ vs FANG
B
Malawi-Alive vs FANG
C
Cameron-Alive vs FANG
D
Uganda-Alive vs FANG
-log10(pvalue)
log2FoldChange

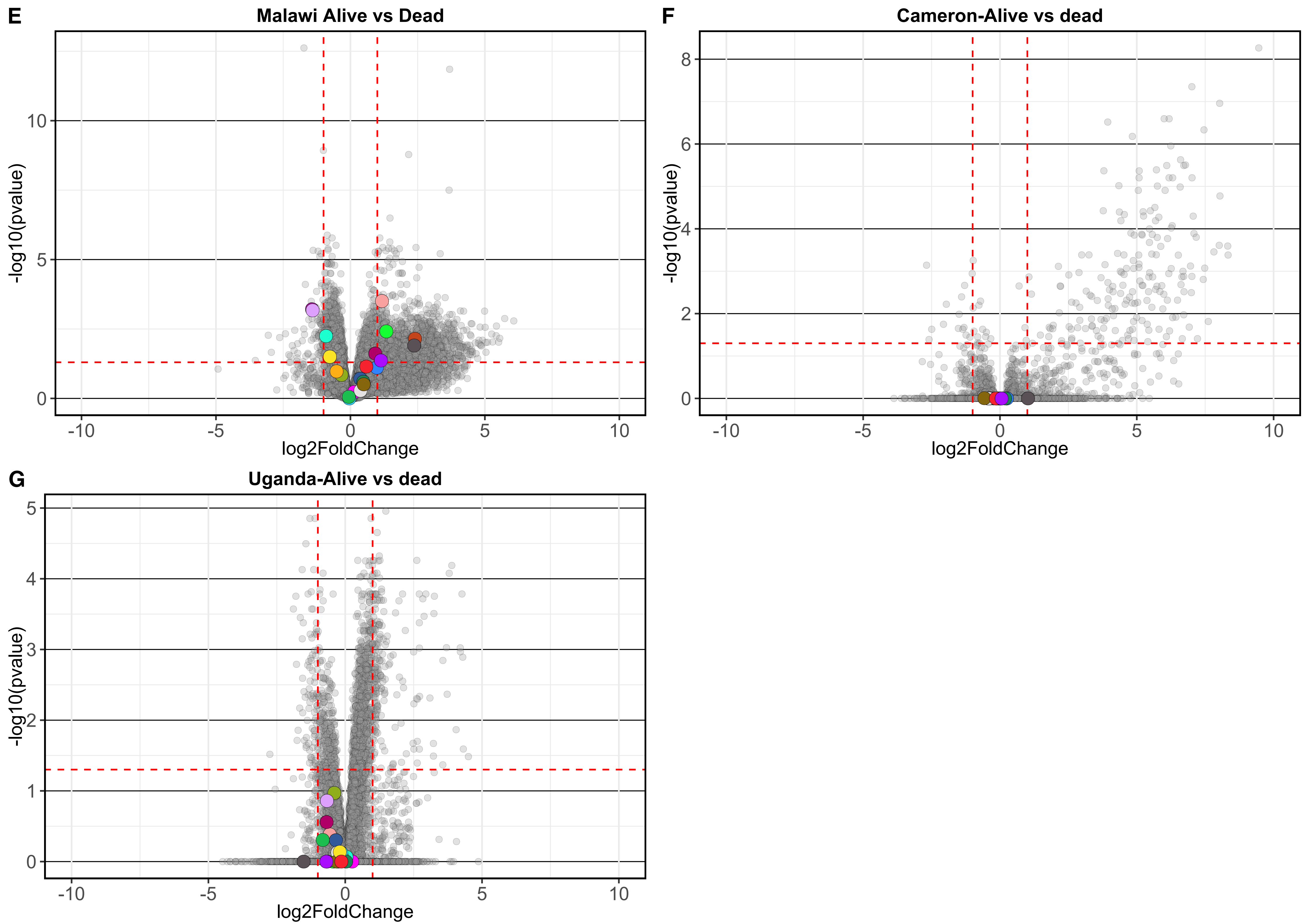
E
Malawi Alive vs Dead
-log10(pvalue)
log2FoldChange
0
5
10
-10
-5
0
5
10
F
Cameron-Alive vs dead
-log10(pvalue)
log2FoldChange
0
2
4
6
8
-10
-5
0
5
10
G
Uganda-Alive vs dead
-log10(pvalue)
log2FoldChange
0
1
2
3
4
5
-10
-5
0
5
10

**Fig. S4. UGTs differential expression in the FUMOZ colony and in field-collected mosquitoes from Malawi, Uganda and Cameroon.** The transcription profiles of the resistant colony FUMOZ, filed-collected resistant mosquitoes from Malawi, Uganda and Cameroon was compared to the FANG transcription profile **(A-D)**. In addition transactional profiles of filed-collected resistant population was compared to the transactional profile of a population of mosquitoes unexposed to permeation from the same  same country **(E-G)**. Differentially regulated genes were defined as those with a corrected p-value threshold of < 0.05 and log2(fold change) > 1 indicated by the dotted red line in each plot. Differentially expressed UGT genes were color coded according to the plot legend.

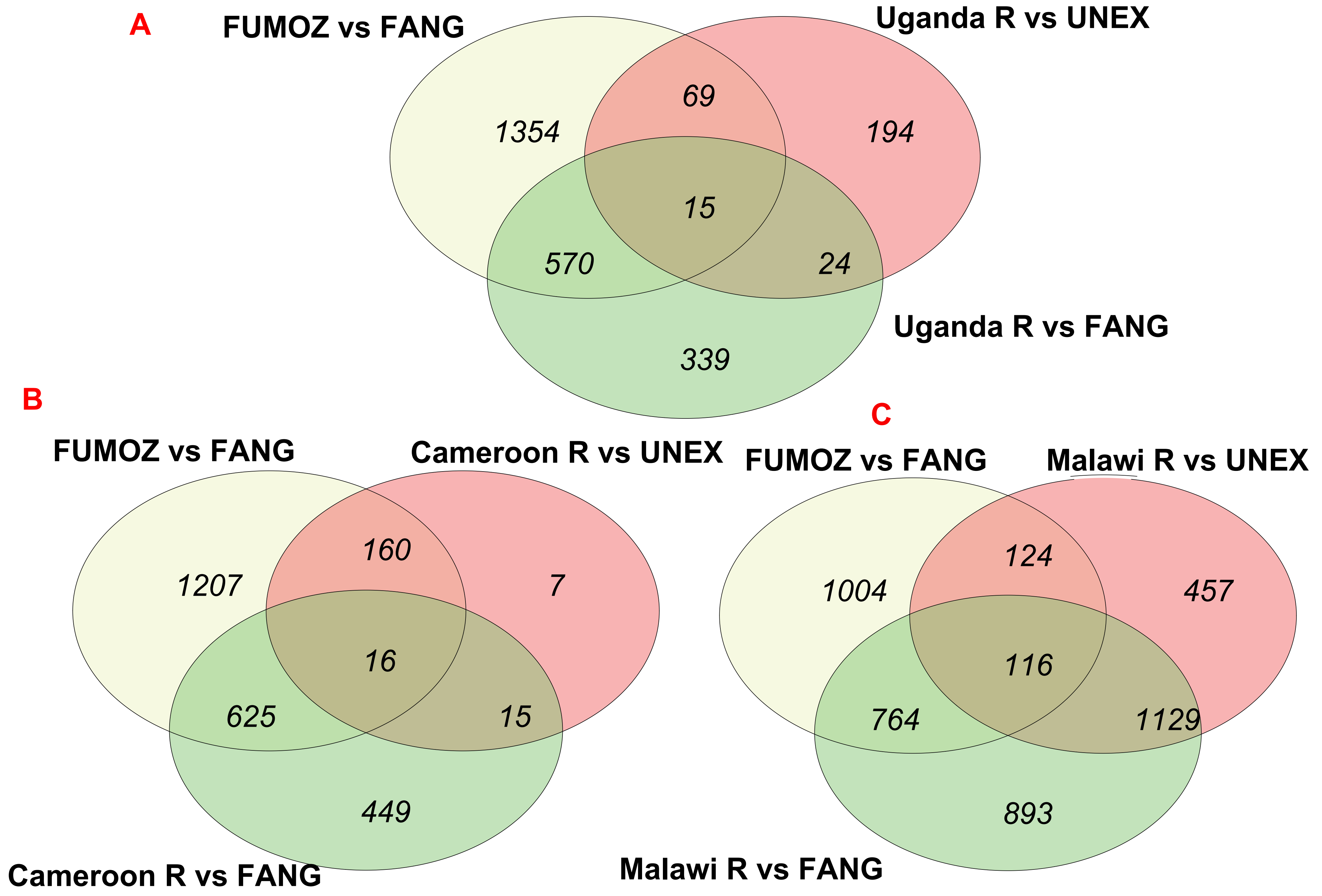
A
FUMOZ vs FANG
Uganda R vs UNEX
Uganda R vs FANG
1354
69
194
15
570
24
339
B
FUMOZ vs FANG
Cameroon R vs UNEX
Cameroon R vs FANG
1207
160
7
16
625
15
449
C
FUMOZ vs FANG
Malawi R vs UNEX
Malawi R vs FANG
1004
124
457
116
764
1129
893

**Fig. S5. Venn diagrams illustrating the comparison of differentially expressed genes.** The diagram displays the number of shared and unique differentially expressed genes between three different scenarios: 1) FUMOZ against FANG, 2) resistant field-collected population (R) against unexposed populations from the same country (UNEX), and 3) resistant field-collected population (R) against FANG. This comparison was conducted in Uganda **(A)**, Cameroon **(B)**, and Malawi **(C)**.

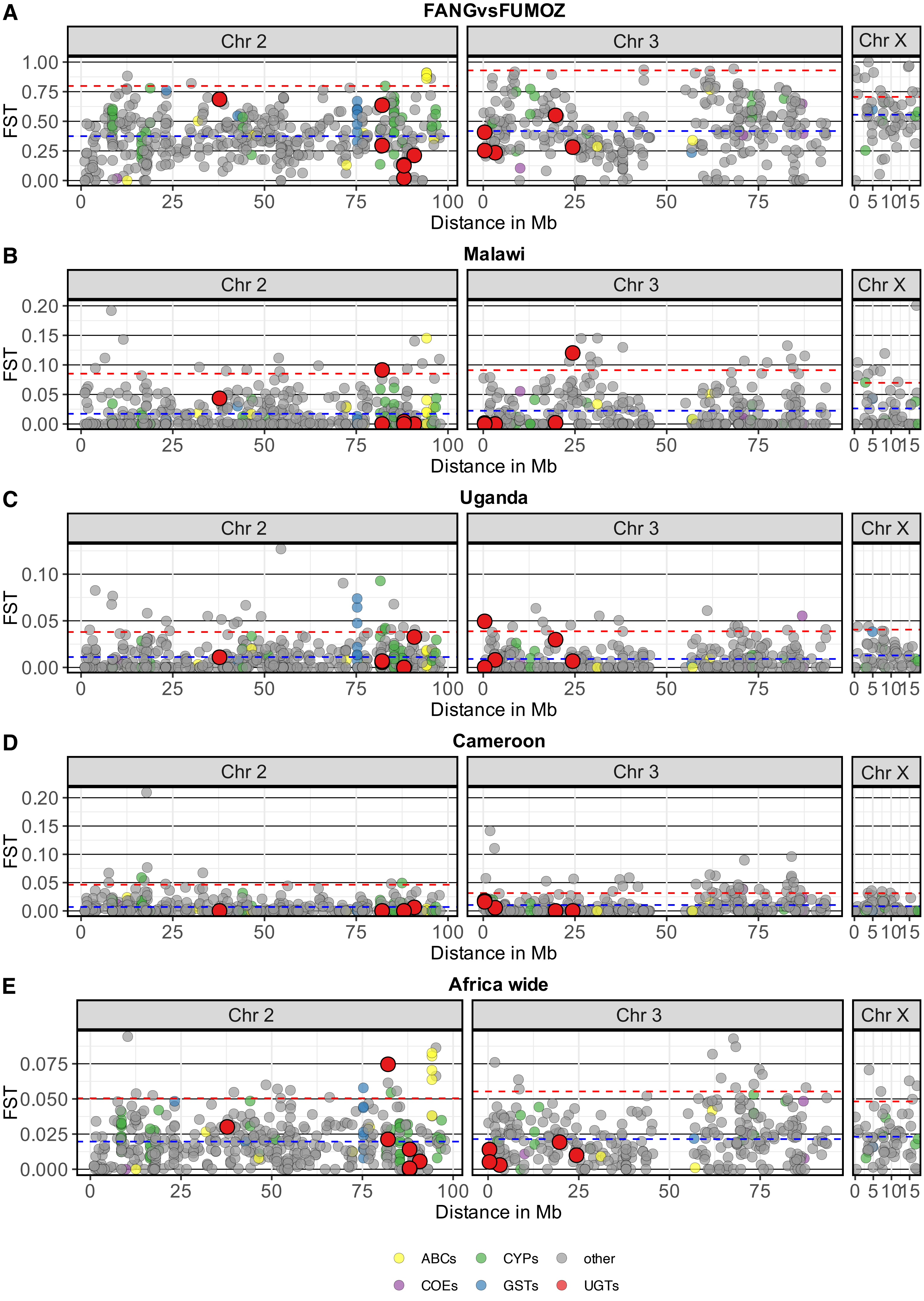
A
FANGvsFUMOZ
B
Malawi
C
Uganda
D
Cameroon
E
Africa wide
Chr 2
Chr 3
Chr X
FST
Distance in Mb
ABCs
CYPs
other
COEs
GSTs
UGTs

**Fig. S6. Gene-wise $F_{st}$ for all genes included in the targeted sequencing in all analyses between resistant and susceptible.** The average and 0.95 quantiles of gene-wise FST for each chromosome are represented by the blue horizontal line and the red horizontal line respectively. The genomic location of all genes in the targeted region (x-axis), represented by a circle, was plotted against the Gene-wise $F_{st}$ (y-axis). Detoxification genes included by the targeted sequencing are color coded according to the plot legend.

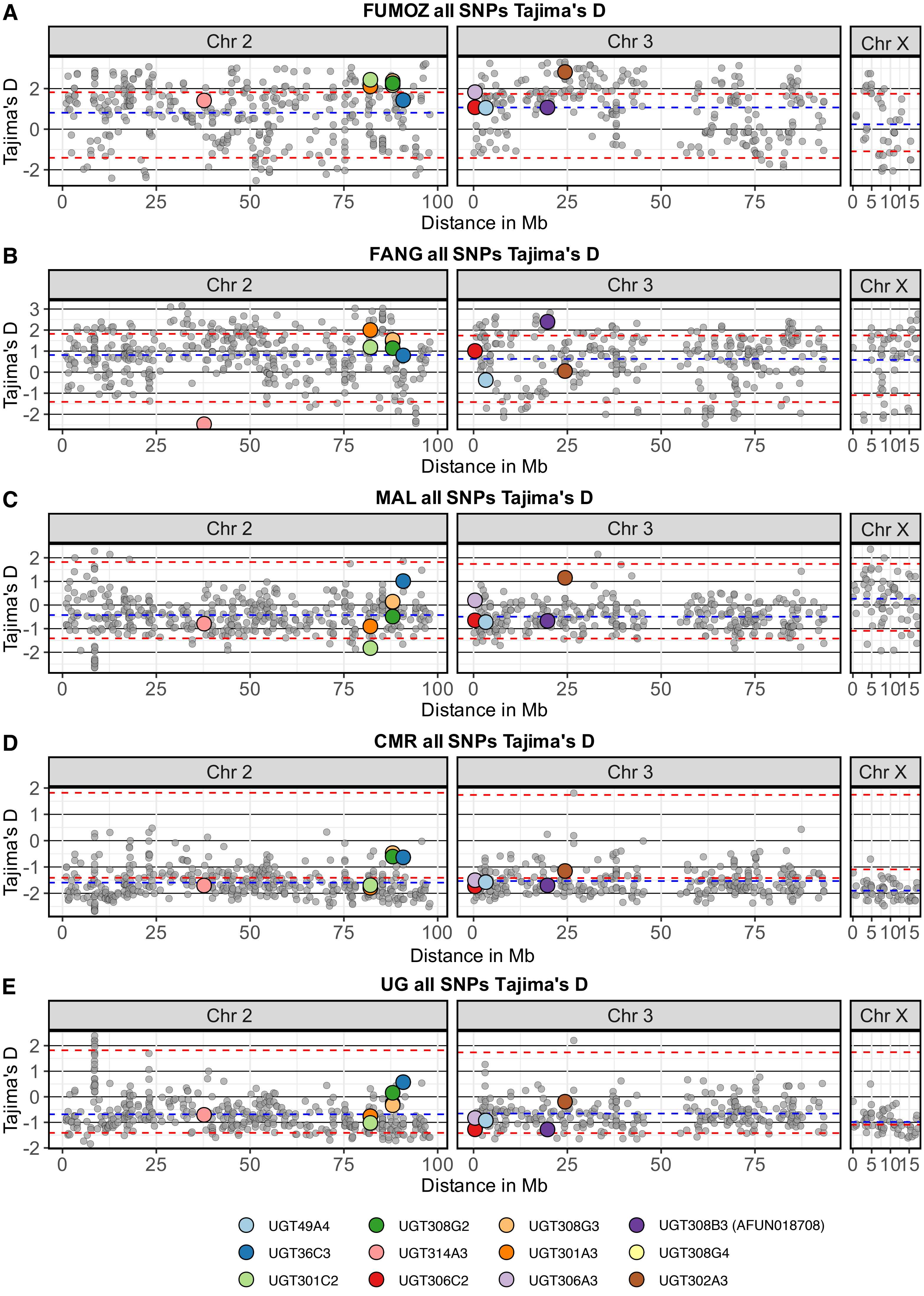
A
FUMOZ all SNPs Tajima's D
B
FANG all SNPs Tajima's D
C
MAL all SNPs Tajima's D
D
CMR all SNPs Tajima's D
E
UG all SNPs Tajima's D
Chr 2
Chr 3
Chr X
Tajima's D
Distance in Mb
UGT49A4
UGT36C3
UGT301C2
UGT308G2
UGT314A3
UGT306C2
UGT308G3
UGT301A3
UGT306A3
UGT308B3 (AFUN018708)
UGT308G4
UGT302A3

**Fig. S7. Gene-wise Tajima's D for all genes included by the targeted enrichment sequencing.** In each country plot, the average is represented by the blue horizontal line and he 0.5 quantiles and 0.95 quantiles are highlighted by the dotted red line. The genomic location of all genes in the targeted region (x-axis), represented by a circle, was plotted against the gene-wise Tajima's D (y-axis). UGT genes included in the targeted sequencing are color coded according to the plot legend.

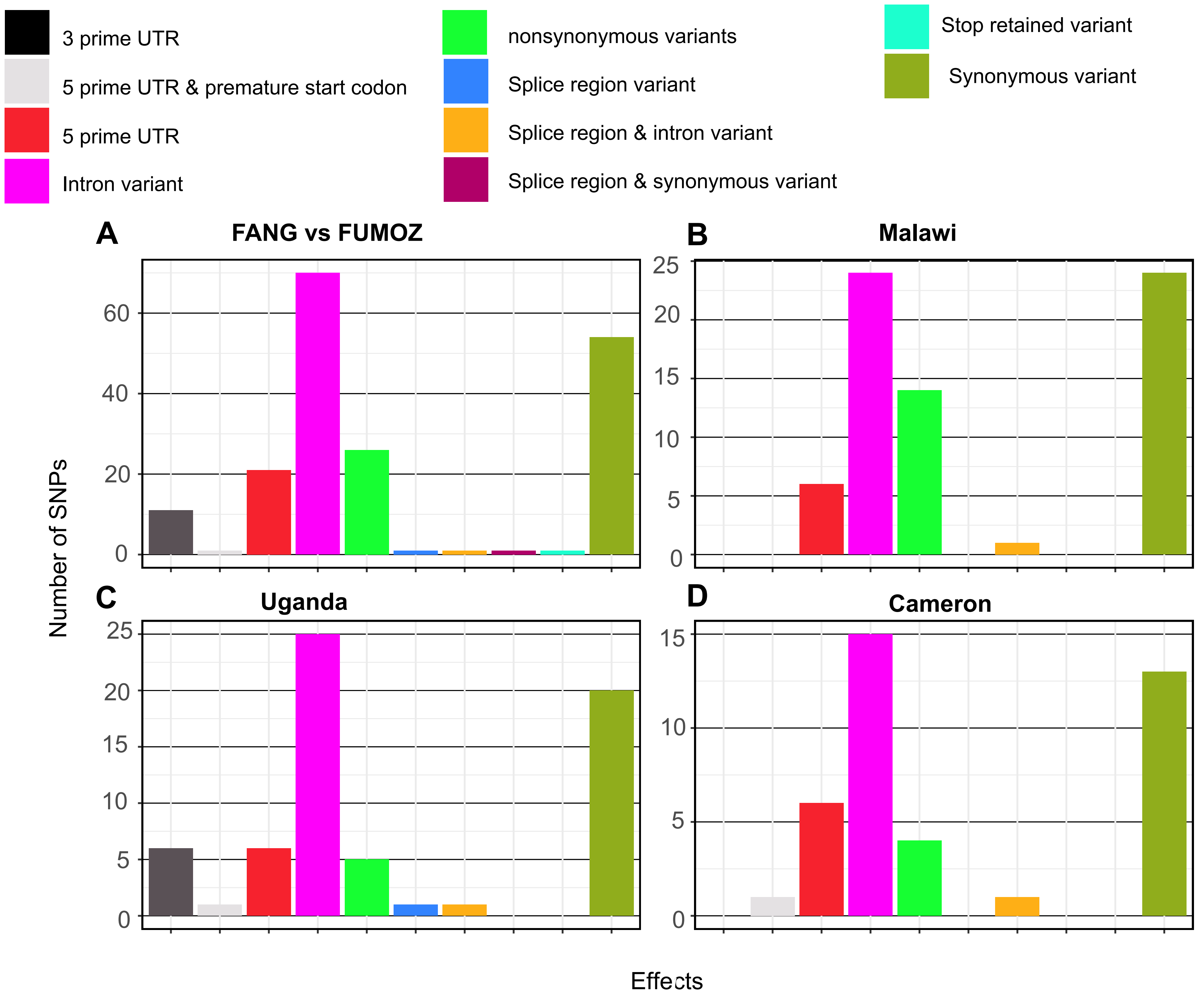

3 prime UTR
5 prime UTR & premature start codon
5 prime UTR
Intron variant
nonsynonymous variants
Splice region variant
Splice region & intron variant
Splice region & synonymous variant
Stop retained variant
Synonymous variant
A
FANG vs FUMOZ
B
Malawi
C
Uganda
D
Cameron
Number of SNPs
Effects

**Fig. S8. Effect caused by UGT genes significantly differentiated SNPs.** Most of the differentiated UGTs' SNPs in all analyses are intronic SNPs. We focused our study on UGT genes non-synonymous SNPs that are significantly differentiated. Details of the number of UGT genes SNPs in each comparison including nonsynonymous SNPs are included in (Table S7).

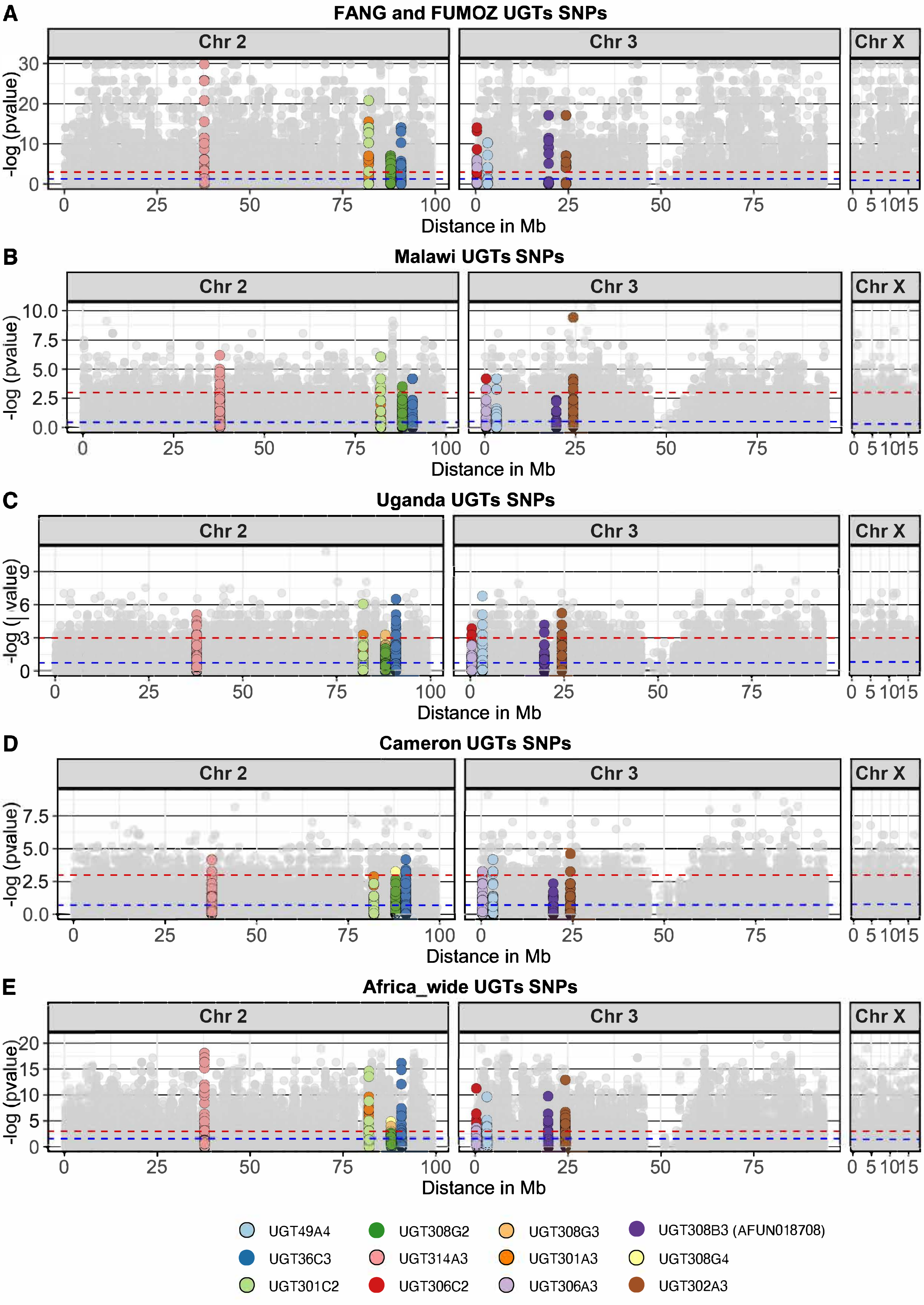

A
FANG and FUMOZ UGTs SNPs
Chr 2
Chr 3
Chr X
-log (pvalue)
Distance in Mb
B
Malawi UGTs SNPs
C
Uganda UGTs SNPs
D
Cameron UGTs SNPs
E
Africa_wide UGTs SNPs
UGT49A4
UGT36C3
UGT301C2
UGT308G2
UGT314A3
UGT306C2
UGT308G3
UGT301A3
UGT306A3
UGT308B3 (AFUN018708)
UGT308G4
UGT302A3

**Fig. S9. UGT genes SNPs divergence between laboratory colonies and in each country.** SNPs located within UGT gene are color-coded according to the plot legend. The blue dotted line indicates the average value in each comparison and the red dotted line indicates the significance level (p-value = 0.05).

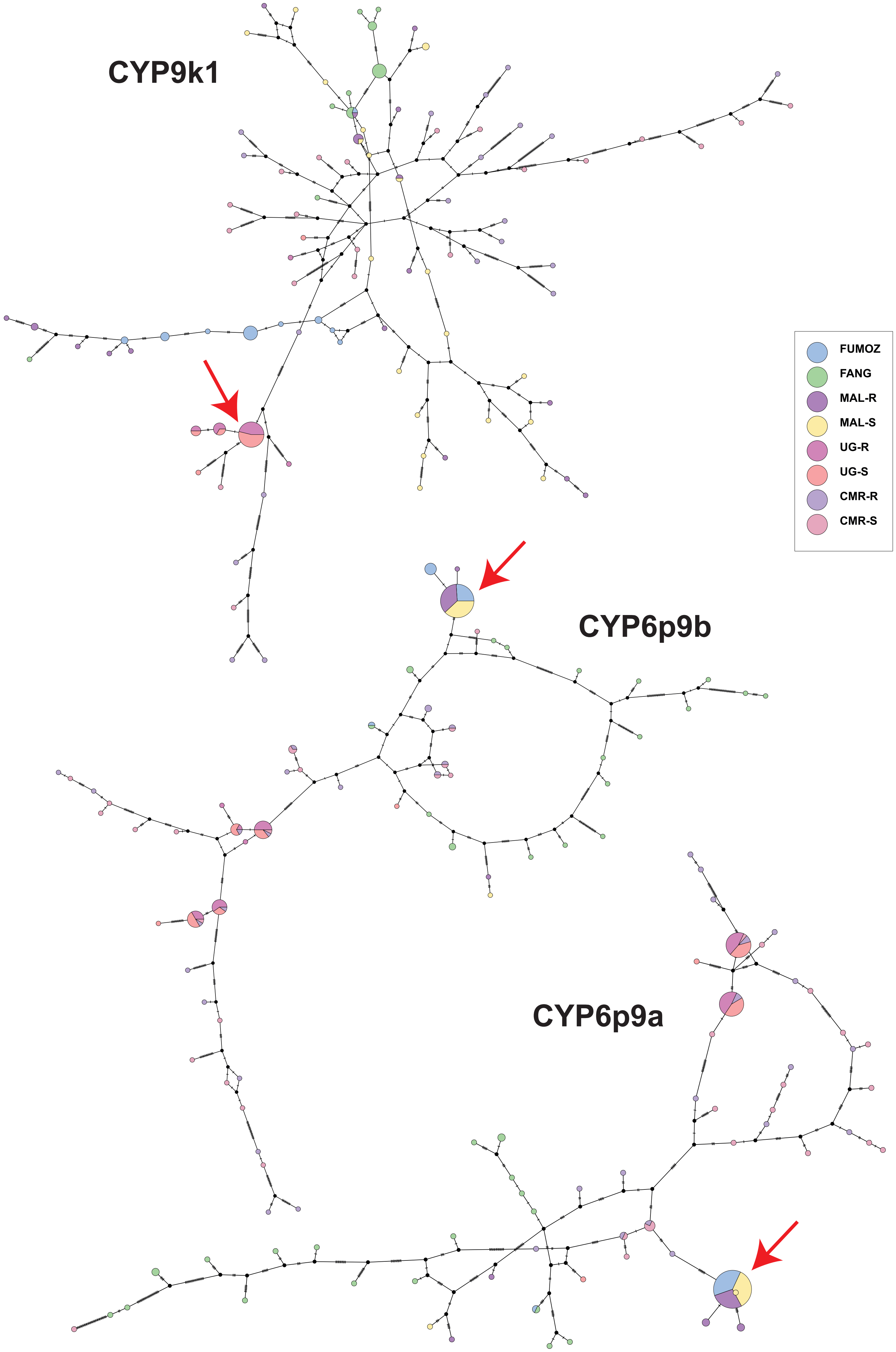
CYP9k1
CYP6p9b
CYP6p9a
FUMOZ
FANG
MAL-R
MAL-S
UG-R
UG-S
CMR-R
CMR-S

**Fig. S10. Hapotype network for *cyp6p9a*, *cyp6p9b* and *cyp9k1*.** TCS haplotype network for *cyp9k1* reveal a directions selection in Uganda supporting previous findings **(A)**. While TCS haplotype network for cyp6p9a and b support previous observation of directional selection in southern Africa represented by Malawi and the resistant FUMOZ colony originally from Mozambique **(B and C)**. The red arrow highlights the dominant haplotype node.

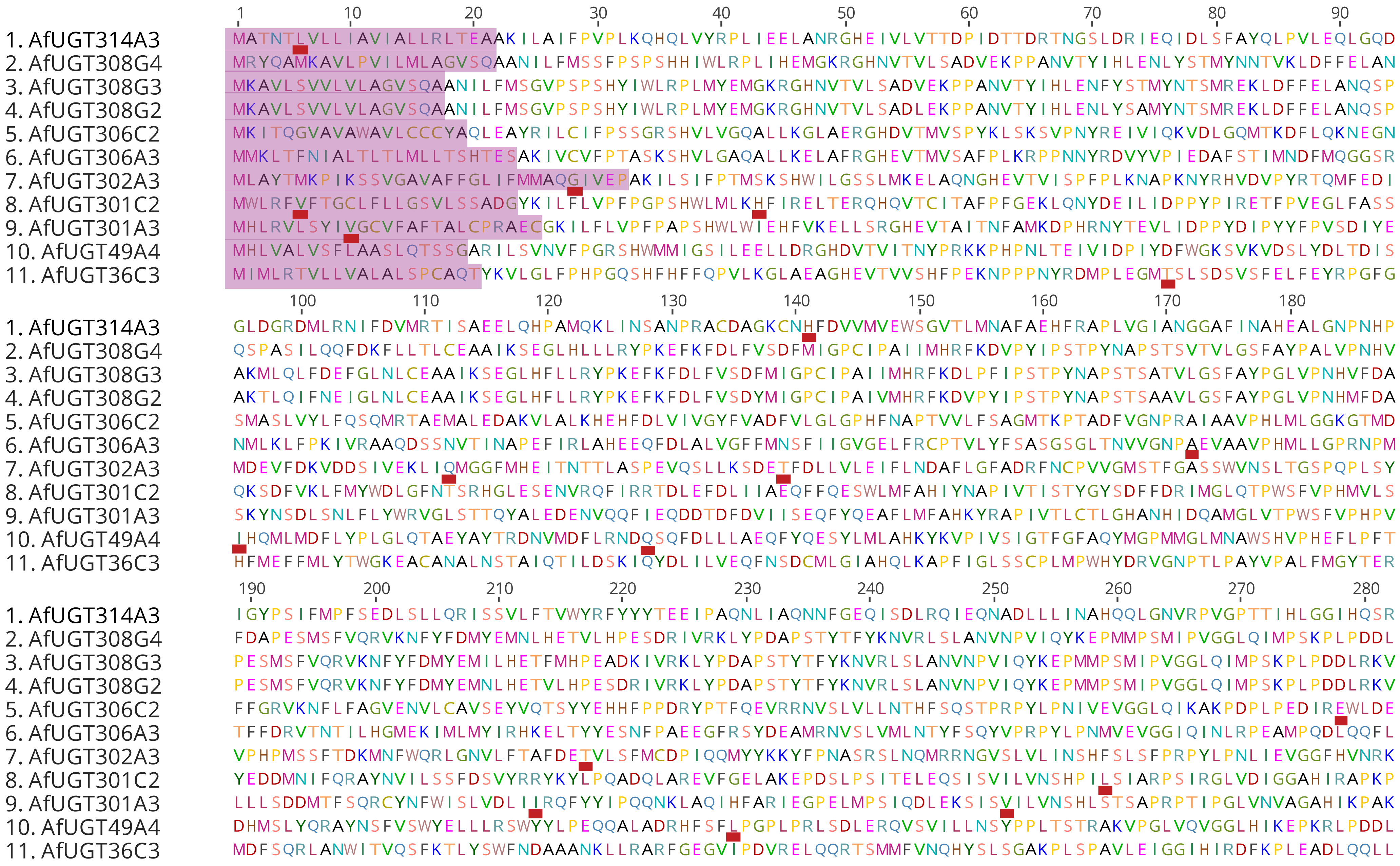
1 10 20 30 40 50 60 70 80 90
1. AfUGT314A3 MATNTLVLLIAVIALLRLTEAAKILAIFPVPLKQHQLVYRPLIEELANRGHEIVLVTTDPIDTTDRTNGSLDRIEQIDLSFAYQLPVLEQLGQD
2. AfUGT308G4 MRYQAMKAVLPVILMLAGVSQAANILFMSSFPSPSHHIWLRPLIHEMGKRGHNVTVLSADVEKPPANVTYIHLENLYSTMYNNTVKLDFFELAN
3. AfUGT308G3 MKAVLSVVLVLAGVSQAANILFMSGVPSPSHYIWLRPLMYEMGKRGHNVTVLSADVEKPPANVTYIHLENFYSTMYNTSMREKLDFFELANQSP
4. AfUGT308G2 MKAVLSVVLVLAGVSQAANILFMSGVPSPSHYIWLRPLMYEMGKRGHNVTVLSADLEKPPANVTYIHLENLYSAMYNTSMREKLDFFELANQSP
5. AfUGT306C2 MKITQGVAVAWAVLCCCYAQLEAYRILCIFPSSGRSHVLVGQALLKGLAERGHDVTMVSPYKLSKSVPNYREIVIQKVDLGQMTKDFLQKNEGN
6. AfUGT306A3 MMKLTFNIALTLTLMLLTSHTESAKIVCVFPTASKSHVLGAQALLKELAFRGHEVTMVSAFPLKRPPNNYRDVYVPIEDAFSTIMNDFMQGGSR
7. AfUGT302A3 MLAYTMKPIKSSVGAVAFFGLIFMMAQGIVEPAKILSIFPTMSKSHWILGSSLMKELAQNGHEVTVISPFPLKNAPKNYRHVDVPYRTQMFEDI
8. AfUGT301C2 MWLRFVFTGCLFLLGSVLSSADGYKILFLVPFPGPSHWLMLKHFIRELTERQHQVTCITAFPFGEKLQNYDEILIDPPYPIRETFPVEGLFASS
9. AfUGT301A3 MHLRVLSYIVGCVFAFTALCPRAECGKILFLVPFPAPSHWLWIEHFVKELLSRGHEVTAITNFAMKDPHRNYTEVLIDPPYDIPYYFPVSDIYE
10. AfUGT49A4 MHLVALVSFLAASLQTSSGARILSVNVFPGRSHWMMIGSILEELLDRGHDVTVITNYPRKKPHPNLTEIVIDPIYDFWGKSVKVDSLYDLTDIS
11. AfUGT36C3 MIMLRTVLLVALALSPCAQTYKVLGLFPHPGQSHFHFFQPVLKGLAEAGHEVTVVSHFPEKNPPPNYRDMPLEGMTSLSDSVSFELFEYRPGFG
100 110 120 130 140 150 160 170 180
1. AfUGT314A3 GLDGRDMLRNIFDVMRTISAEELQHPAMQKLINSANPRACDAGKCNHFDVVMVEWSGVTLMNAFAEHFRAPLVGIANGGAFINAHEALGNPNHP
2. AfUGT308G4 QSPASILQQFDKFLLTLCEAAIKSEGLHLLLRYPKEFKFDLFVSDFMIGPCIPAIIMHRFKDVPYIPSTPYNAPSTSVTVLGSFAYPALVPNHV
3. AfUGT308G3 AKMLQLFDEFGLNLCEAAIKSEGLHFLLRYPKEFKFDLFVSDFMIGPCIPAIIMHRFKDLPFIPSTPYNAPSTSATVLGSFAYPGLVPNHVFDA
4. AfUGT308G2 AKTLQIFNEIGLNLCEAAIKSEGLHFLLRYPKEFKFDLFVSDYMIGPCIPAIVMHRFKDVPYIPSTPYNAPSTSAAVLGSFAYPGLVPNHMFDA
5. AfUGT306C2 SMASLVYLFQSQMRTAEMALEDAKVLALKHEHFDLVIVGYFVADFVLGLGPHFNAPTVVLFSAGMTKPTADFVGNPRAIAAVPHLMLGGKGTMD
6. AfUGT306A3 NMLKLFPKIVRAAQDSSNVTINAPEFIRLAHEEQFDLALVGFFMNSFIIGVGELFRCPTVLYFSASGSGLTNVVGNPAEVAAVPHMLLGPRNPM
7. AfUGT302A3 MDEVFDKVDDSIVEKLIQMGGFMHEITNTTLASPEVQSLLKSDETFDLLVLEIFLNDAFLGFADRFNCPVVGMSTFGASSWVNSLTGSPQPLSY
8. AfUGT301C2 QKSDFVKLFMYWDLGFNTSRHGLESENVRQFIRRTDLEFDLIIAEQFFQESWLMFAHIYNAPIVTISTYGYSDFFDRIMGLQTPWSFVPHMVLS
9. AfUGT301A3 SKYNSDLSNLFLYWRVGLSTTQYALEDENVQQFIEQDDTDFDVIISEQFYQEAFLMFAHKYRAPIVTLCTLGHANHIDQAMGLVTPWSFVPHPV
10. AfUGT49A4 LHQMLMDFLYPLGLQTAEYAYTRDNVMDFLRNDQSQFDLLLAEQFYQESYLMLAHKYKVPIVSIGTFGFAQYMGPMMGLMNAWSHVPHEFLPFT
11. AfUGT36C3 HFMEFFMLYTWGKEACANALNSTAIQTILDSKIQYDLILVEQFNSDCMLGIAHQLKAPFIGLSSCPLMPWHYDRVGNPTLPAYVPALFMGYTER
190 200 210 220 230 240 250 260 270 280
1. AfUGT314A3 IGYPSIFMPFSEDLSLLQRISSVLFTVWYRFYYYTEEIPAQNLIAQNNFGEQISDLRQIEQNADLLLINAHQQLGNVRPVGPTTIHLGGIHQSR
2. AfUGT308G4 FDAPESMSFVQRVKNFYFDMYEMNLHETVLHPESDRIVRKLYPDAPSTYTFYKNVRLSLANVNPVIQYKEPMMPSMIPVGGLQIMPSKPLPDDL
3. AfUGT308G3 PESMSFVQRVKNFYFDMYEMILHETFMHPEADKIVRKLYPDAPSTYTFYKNVRLSLANVNPVIQYKEPMMPSMIPVGGLQIMPSKPLPDDLRKV
4. AfUGT308G2 PESMSFVQRVKNFYFDMYEMNLHETVLHPESDRIVRKLYPDAPSTYTFYKNVRLSLANVNPVIQYKEPMMPSMIPVGGLQIMPSKPLPDDLRKV
5. AfUGT306C2 FFGRVKNFLFAGVENVLCAVSEYVQTSYYEHHFPPDRYPTFQEVRRNVSLVLLNTHFSQSTPRPYLPNIVEVGGLQIKAKPDPLPEDIREWLDE
6. AfUGT306A3 TFFDRVTNTILHGMEKIMLMYIRHKELTYYESNFPAEEGFRSYDEAMRNVSLVMLNTYFSQYVPRPYLPNMVEVGGIQINLRPEAMPQDLQQFL
7. AfUGT302A3 VPHPMSSFTDKMNFWQRLGNVLFTAFDETVLSFMCDPIQQMYYKKYFPNASRSLNQMRRNGVSLVLINSHFSLSFPRPYLPNLIEVGGFHVNRK
8. AfUGT301C2 YEDDMNIFQRAYNVILSSFDSVYRRYKYLPQADQLAREVFGELAKEPDSLPSITELEQSISVILVNSHPILSIARPSIRGLVDIGGAHIRAPKP
9. AfUGT301A3 LLLSDDMTFSQRCYNFWISLVDLIIRQFYYIPQQNKLAQIHFARIEGPELMPSIQDLEKSISVILVNSHLSTSAPRPTIPGLVNVAGAHIKPAK
10. AfUGT49A4 DHMSLYQRAYNSFVSWYELLLRSWYYLPEQQALADRHFSFLPGPLPRLSDLERQVSVILLNSYPPLTSTRAKVPGLVQVGGLHIKEPKRLPDDL
11. AfUGT36C3 MDFSQRLANWITVQSFKTLYSWFNDAAANKLLRARFGEGVIPDVRELQQRTSMMFVNQHYSLSGAKPLSPAVLEIGGIHIRDFKPLEADLQQLL

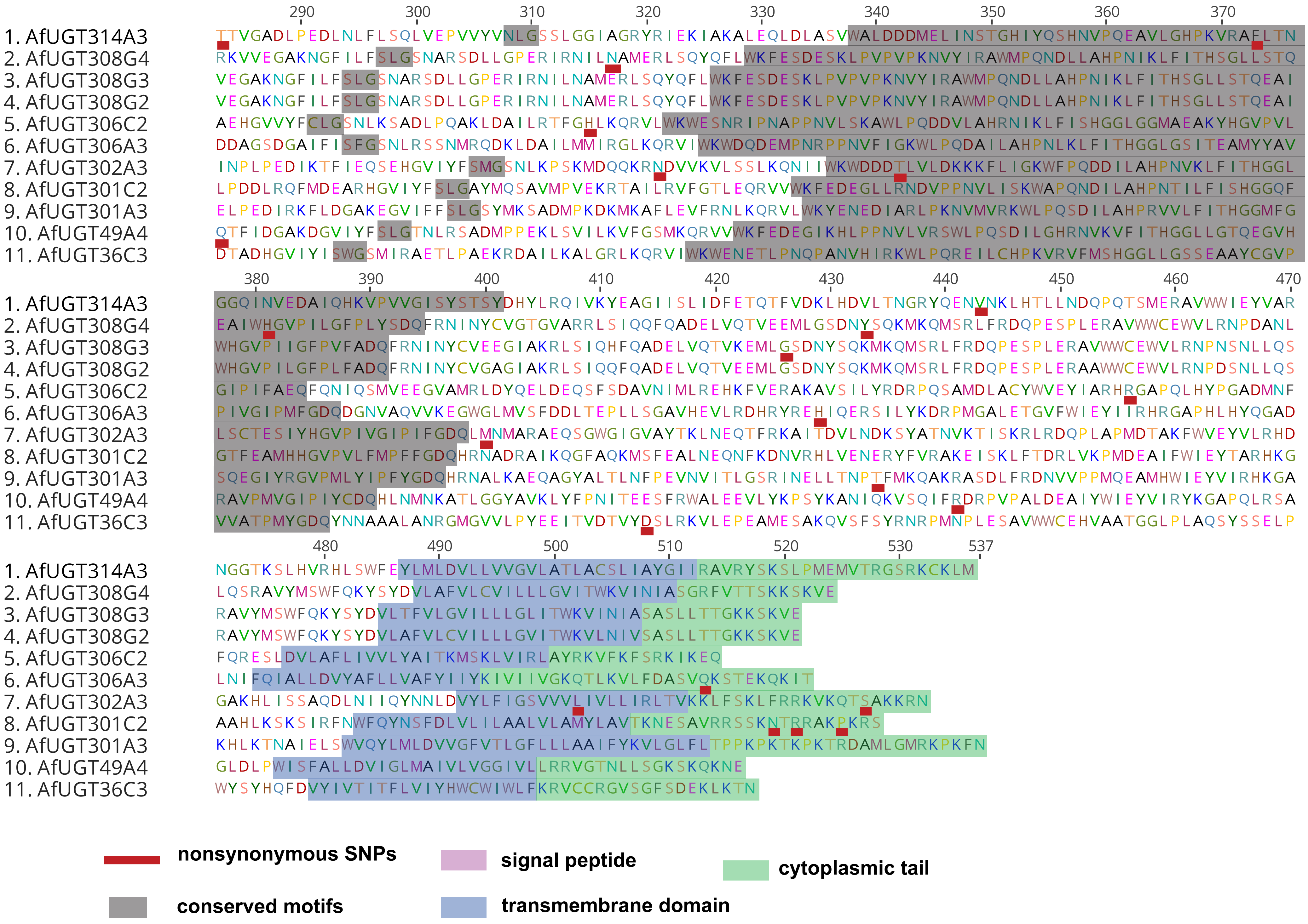

1. AfUGT314A3
2. AfUGT308G4
3. AfUGT308G3
4. AfUGT308G2
5. AfUGT306C2
6. AfUGT306A3
7. AfUGT302A3
8. AfUGT301C2
9. AfUGT301A3
10. AfUGT49A4
11. AfUGT36C3
nonsynonymous SNPs
signal peptide
cytoplasmic tail
conserved motifs
transmembrane domain

**Fig. S11. nonsynonymous SNPs location in UGTs targeted by sequencing.** Functional protein regions in non-aligned UGTs included in the targeted enrichment sequencing are highlighted in the figure legend. Nonsynonymus SNPs that are significantly differentiated between putatively susceptible and resistant from Malawi, Cameroon, and Uganda, and between the FANG and the FUMOZ colony are highlighted with a red color in the protein sequence. For further details on those SNPs see Table S8.
