## Supplementary Tables for "Overexpression and nonsynonymous mutations of UDP-glycosyltransferases potentially associated with pyrethroid resistance in *Anopheles funestus*"

**Supplementary text 1. Significant upregulation of P450 genes in resistant populations was detected confirming previous findings.**

Significant upregulation of Phase I cytochrome P450 enzymes, compared to the UGTs, was detected when transcriptional profiling of permethrin-resistant populations was compared to that of susceptible lab colony (FANG) confirming previously published observations (Supplemental Figure 3) [17, 26]. A significant upregulation in *CYP6P9a* and *CYP6P9b* in all cohorts was detected, in the FUMOZ and in the resistant population from Malawi they were the highest expressed P450 enzymes, and significantly upregulated in the resistant population from Cameroon and Uganda. Roles of both genes in pyrethroid resistance were previously characterised [1, 2], and their role in pyrethroid metabolism and reduction in bed nets efficacy was investigated [3-6]. Other P450 enzymes were highly overexpressed in mosquitoes collected from other regions, *CYP325A* (AFUN015966) and *CYP6P5* (AFUN015888) are highly expressed in Cameroon, CYP9K1 (AFUN007549) and CYP6P5 (AFUN015888) were detected to be highly expressed in Uganda, while CYP9J11 is expressed universally in all investigated countries across Africa [5, 7, 8]. Other Phase II and Phase III enzymes have been detected in resistant populations across all cohorts notably an ABC transporter (AFUN019220) and a *GSTe2* (AFUN015809). The role of polymorphism and overexpression of *GSTe2* in conferring high-level resistance to DDT and cross-resistance to pyrethroid has been previously highlighted (Supplementary Fig. 3, Supplementary Dataset 4) [9].

**Supplementary text 2. VectorBase Gene IDs from FUMOZ strain reference genome annotation of detoxification genes targeted by the enrichment.**

**12 genes from UDP-glucuronyl transferases (UGTs) family:**

"AFUN005498" "AFUN016158" "AFUN004354" "AFUN019724" "AFUN009064" "AFUN004976" "AFUN003593" "AFUN005786" "AFUN016302" "AFUN002058" "AFUN018708" "AFUN019845"

**14 genes ABC transporters (ABCs):**

"AFUN019220" "AFUN008521" "AFUN020102" "AFUN008943" "AFUN003077" "AFUN015982" "AFUN015980" "AFUN015978" "AFUN015745" "AFUN005260" "AFUN010004" "AFUN016307" "AFUN016414" "AFUN007162"

**5 genes carboxylesterases (COEs):**

"AFUN007734" "AFUN000373" "AFUN010202" "AFUN002517" "AFUN002514"

**66 genes cytochrome P450 monooxygenases:**

"AFUN015786" "AFUN015785" "AFUN008357" "AFUN015792" "AFUN015889" "AFUN015888" "AFUN020895" "AFUN019365" "AFUN015802" "AFUN015801" "AFUN015714" "AFUN005865" "AFUN005864" "AFUN015993" "AFUN015992" "AFUN015990" "AFUN015991" "AFUN004720" "AFUN015790" "AFUN015791" "AFUN015957" "AFUN015956" "AFUN003690" "AFUN002408" "AFUN004312" "AFUN004315" "AFUN004316" "AFUN015918" "AFUN015919" "AFUN019401" "AFUN015795" "AFUN015794" "AFUN010918" "AFUN020636" "AFUN015961" "AFUN015960" "AFUN015962" "AFUN015963" "AFUN015797" "AFUN015796" "AFUN015776" "AFUN015775" "AFUN015774" "AFUN016005" "AFUN015938" "AFUN020179" "AFUN006135" "AFUN006122" "AFUN015723" "AFUN004884" "AFUN015908" "AFUN015904" "AFUN015906" "AFUN004597" "AFUN002275" "AFUN000518" "AFUN001382" "AFUN001383" "AFUN015865" "AFUN015866" "AFUN006702" "AFUN011481" "AFUN005715" "AFUN007549" "AFUN010428" "AFUN021097"

**12 genes glutathione S-transferases (GSTs):**

"AFUN008426" "AFUN011410" "AFUN015715" "AFUN016008" "AFUN015811" "AFUN015810" "AFUN015809" "AFUN015807" "AFUN001774" "AFUN015808" "AFUN008829" "AFUN007291"

**Other genes 698**

**Table S1. The number of bi-allelic sites per analysis per chromosome detected using popgenome R package for gene-wise AFS statistics.**

| **Chromosomes** | **Chr2** | **Chr3** | **ChrX** | **Total** |
| --- | --- | --- | --- | --- |
| **Genes targeted** | 431 | 315 | 61 | 807 |
| **FUMOZ and FANG** | 16,623 | 10,332 | 1,611 | 28,566 |
| **Malawi** | 27,132 | 20,130 | 2,500 | 49,762 |
| **Cameron** | 50,652 | 33,335 | 7,632 | 91,619 |
| **Uganda** | 48,969 | 33,494 | 7,877 | 90,340 |
| **Africa-wide** | 77,196 | 47,573 | 11,579 | 136,348 |

**Table S2. Average gene-wise *F_ST_* values and *F_ST_* values at the 0.8 and 0.95 quantiles in each analysis.**

| **Analysis** | **Fst average and 0.8 quantile** | **Chr2** | **Chr3** | **ChrX** |
| --- | --- | --- | --- | --- |
| **Africa_wide** | Average | 0.020 | 0.021 | 0.023 |
|  | 0.8 quantile | 0.031 | 0.035 | 0.032 |
|  | 0.95 quantile | 0.048 | 0.051 | 0.055 |
| **FANGvsFUMOZ** | Average | 0.375 | 0.418 | 0.555 |
|  | 0.8 quantile | 0.527 | 0.620 | 0.695 |
|  | 0.95 quantile | 0.706 | 0.798 | 0.929 |
| **Malawi** | Average | 0.017 | 0.022 | 0.026 |
|  | 0.8 quantile | 0.032 | 0.045 | 0.048 |
|  | 0.95 quantile | 0.069 | 0.085 | 0.091 |
| **Uganda** | Average | 0.011 | 0.009 | 0.013 |
|  | 0.8 quantile | 0.020 | 0.017 | 0.023 |
|  | 0.95 quantile | 0.041 | 0.038 | 0.039 |
| **Cameron** | Average | 0.007 | 0.010 | 0.008 |
|  | 0.8 quantile | 0.012 | 0.018 | 0.017 |
|  | 0.95 quantile | 0.031 | 0.046 | 0.032 |

**Table S3. Gene-wise *F_ST_* values for UGT genes that were included in the target enrichment analysis.** Highlighted cells indicate values above 0.95 (red) and 0.8 (yellow) quantiles of gene-wise *F_ST_* values for the respective chromosomes.

| **gene_ID** | **UGT_name** | **Chromosome** | **Africa wide** | **FANG and FUMOZ** | **Malawi** | **Uganda** | **Cameron** |
| --- | --- | --- | --- | --- | --- | --- | --- |
| **AFUN002058** | UGT49A4 | chr3 | 0.00298 | 0.23497 | 0 | 0.00814 | 0.00555 |
| **AFUN003593** | UGT36C3 | chr2 | 0.00563 | 0.21253 | 0 | 0.03263 | 0.00623 |
| **AFUN004354** | UGT301C2 | chr2 | 0.07463 | 0.63575 | 0.09142 | 0.00555 | 0 |
| **AFUN004976** | UGT308G2 | chr2 | 0.01416 | 0.13249 | 0.00417 | 0 | 0 |
| **AFUN005498** | UGT314A3 | chr2 | 0.03007 | 0.68614 | 0.04346 | 0.01091 | 0 |
| **AFUN005786** | UGT306C2 | chr3 | 0.01398 | 0.40737 | 0.00165 | 0.0494 | 0.01811 |
| **AFUN009064** | UGT308G3 | chr2 | 0 | 0.02129 | 0 | 0 | 0 |
| **AFUN016158** | UGT301A3 | chr2 | 0.02123 | 0.29558 | 0 | 0.00685 | 0 |
| **AFUN016302** | UGT306A3 | chr3 | 0.00521 | 0.25304 | 0 | 0 | 0.01606 |
| **AFUN018708** | *not assigned* | chr3 | 0.01929 | 0.5499 | 0.00211 | 0.02985 | 0 |
| **AFUN019724** | UGT308G4 | chr2 | 0.00068 | 0.12529 | 0 | 0 | 0 |
| **AFUN019845** | UGT302A3 | chr3 | 0.00997 | 0.28193 | 0.12021 | 0.00678 | 0 |

**Table S4. Average and median Tajima’s D values per chromosome in each population compared to the coalescent simulation.**

| **Average** | **Populations** | **Chr2** | **Chr3** | **ChrX** |
| --- | --- | --- | --- | --- |
|  | FANG | 0.8143 | 0.6291 | 0.5813 |
|  | FUMOZ | 0.8102 | 1.0696 | 0.2392 |
|  | Malawi | -0.4246 | -0.4953 | 0.2645 |
|  | Uganda | -0.6846 | -0.6552 | -0.9853 |
|  | Cameron | -1.5948 | -1.5348 | -1.8989 |
| **Median** | FANG | 0.8670 | 0.8482 | 0.9221 |
|  | FUMOZ | 1.0689 | 1.3313 | 0.0102 |
|  | Malawi | -0.4455 | -0.5343 | 0.3631 |
|  | Uganda | -0.8283 | -0.7323 | -1.0907 |
|  | Cameron | -1.7004 | -1.6584 | -1.8885 |
| **Coalescent simulation** | | | | |
| Average | | 0.2218 | 0.0958 | 0.2379 |
| Median | | 0.2173 | 0.0405 | 0.0102 |
| 0.05 quantile | | -1.4109 | -1.4201 | -1.0922 |
| 0.2 quantile | | -0.5571 | -0.7993 | -0.6209 |
| 0.8 quantile | | 1.0212 | 1.0503 | 0.9700 |
| 0.95 quantile | | 1.8189 | 1.7375 | 1.7443 |

**Table S5. Gene-wise Tajima’s D values for targeted UGT genes by the enrichment analysis.** Highlighted cells in green are below 0.05 quantiles of simulated Tajima’s D values and highlighted cells in yellow are above 0.95 quantiles of simulated Tajima’s D values.

| **UGT genes** | **UGT_name** | **Chromosome** | **FANG** | **FUMOZ** | **Malawi** | **Uganda** | **Cameron** |
| --- | --- | --- | --- | --- | --- | --- | --- |
| **AFUN002058** | UGT49A4 | chr3 | -0.3728 | 1.0542 | -0.7221 | -0.9265 | -1.5768 |
| **AFUN003593** | UGT36C3 | chr2 | 0.7954 | 1.428 | 1.013 | 0.5714 | -0.6343 |
| **AFUN004354** | UGT301C2 | chr2 | 1.1838 | 2.4351 | -1.8243 | -1.0221 | -1.6931 |
| **AFUN004976** | UGT308G2 | chr2 | 1.1402 | 2.2638 | -0.4893 | 0.1556 | -0.6061 |
| **AFUN005498** | UGT314A3 | chr2 | -2.4709 | 1.415 | -0.7884 | -0.7068 | -1.7034 |
| **AFUN005786** | UGT306C2 | chr3 | 1.0094 | 1.0781 | -0.6474 | -1.2729 | -1.7345 |
| **AFUN009064** | UGT308G3 | chr2 | 1.5294 | 2.3747 | 0.142 | -0.3293 | -0.4731 |
| **AFUN016158** | UGT301A3 | chr2 | 2.0008 | 2.1127 | -0.9027 | -0.7618 | -1.7911 |
| **AFUN016302** | UGT306A3 | chr3 | NA | 1.8265 | 0.1978 | -0.8295 | -1.5029 |
| **AFUN018708** | *not assigned* | chr3 | 2.3948 | 1.0695 | -0.6716 | -1.2779 | -1.6972 |
| **AFUN019724** | UGT308G4 | chr2 | 1.446 | 2.169 | 0.1264 | -0.2672 | -0.5817 |
| **AFUN019845** | UGT302A3 | chr3 | 0.0534 | 2.8176 | 1.1518 | -0.1929 | -1.1523 |

**Table S6. Number of SNPs retained in each comparison for *F_ST_* -based analysis within the region targeted by enrichment sequencing.**

| **Analysis** | **Total number of SNPs** | **Number of SNPs with significant pFst** | **Number of effects caused by SNPs with significant pFst** | **Significant non-synonymous SNPs** |
| --- | --- | --- | --- | --- |
| **Africa-wide** | 136,416 | 13,653 | 43,771 | 1791 |
| **FANG and FUMOZ** | 149,170 | 10,414 | 33,129 | 1275 |
| **Malawi** | 156,843 | 3,088 | 9,432 | 406 |
| **Cameron** | 157,698 | 2,538 | 7,975 | 264 |
| **Uganda** | 153,228 | 3,047 | 9,505 | 304 |

**Table S7. Number of SNPs associated with UGT genes**

| Total number of | SNPs | Total  nonsynonymous variants | SNPs with significant pFst | Synonymous variants | Non-synonymous variants | SNPs on the 5 prime UTR | SNPs on the 3 prime UTR | Intron SNPs | Splice regions SNPs |
| --- | --- | --- | --- | --- | --- | --- | --- | --- | --- |
| Africa-wide | 1470 | 311 | 546 | 70 | 32 | 27 | 9 | 82 | 0 |
| FANG and FUMOZ | 1496 | 314 | 456 | 54 | 26 | 21 | 11 | 70 | 1 |
| Malawi | 1501 | 314 | 174 | 24 | 14 | 6 | 0 | 24 | 0 |
| Cameron | 1511 | 314 | 81 | 13 | 4 | 6 | 0 | 15 | 0 |
| Uganda | 1516 | 311 | 131 | 20 | 5 | 6 | 6 | 25 | 1 |

**Table S8. Significantly divergent non-synonymous SNPs in all countries.**

| **Pos** | **pfst** | **qvalue** | **Alive** | **Dead** | **wcFst** | **R** | **A** | **Gene ID** | **UGT_Name** | **C_change** | **A_A_change** | **Population** |
| --- | --- | --- | --- | --- | --- | --- | --- | --- | --- | --- | --- | --- |
| 3258498 | 0.0153301 | 0.888644509 | 0 | 0.2 | 0.166667 | A | G | AFUN002058 | UGT49A4 | c.383A>G | p.Gln128Arg | Cameroon Dead & Alive |
| 24377219 | 0.038206 | 0.888644509 | 0.15 | 0 | 0.111111 | G | C | AFUN019845 | UGT302A3 | c.14C>G | p.Thr5Ser | Cameroon Dead & Alive |
| 37739309 | 0.0381976 | 0.888644509 | 0 | 0.15 | 0.111111 | G | A | AFUN005498 | UGT314A3 | c.1327G>A | p.Val443Ile | Cameroon Dead & Alive |
| 90805096 | 0.0442079 | 0.8886445 | 0.8 | 0.5 | 0.133333 | A | T | AFUN003593 | UGT36C3 | c.1242T>A | p.Asp414Glu | Cameroon Dead & Alive |
| 382438 | 7.80E-07 | 3.80E-05 | 0 | 0.65 | 0.615385 | T | A | AFUN005786 | UGT306C2 | c.833A>T | p.Glu278Val | FUMOZ & FANG |
| 384197 | 0.00234011 | 0.035273515 | 0.7 | 1 | 0.277778 | G | C | AFUN016302 | UGT306A3 | c.1287C>G | p.His429Gln | FUMOZ & FANG |
| 385026 | 0.00327112 | 0.047143634 | 0.3 | 0 | 0.277778 | G | A | AFUN016302 | UGT306A3 | c.515C>T | p.Ala172Val | FUMOZ & FANG |
| 3259511 | 3.49E-05 | 0.000997223 | 0.5 | 0 | 0.5 | G | T | AFUN002058 | UGT49A4 | c.1322G>T | p.Arg441Leu | FUMOZ & FANG |
| 24375373 | 0.000875012 | 0.015288292 | 0.35 | 0 | 0.333333 | C | T | AFUN019845 | UGT302A3 | c.1580G>A | p.Ser527Asn | FUMOZ & FANG |
| 24375837 | 0.000874524 | 0.015288292 | 0.35 | 0 | 0.333333 | T | A | AFUN019845 | UGT302A3 | c.1198A>T | p.Met400Leu | FUMOZ & FANG |
| 24376073 | 0.000874087 | 0.015288292 | 0.35 | 0 | 0.333333 | T | C | AFUN019845 | UGT302A3 | c.962A>G | p.Asn321Ser | FUMOZ & FANG |
| 37735174 | 0.00234051 | 0.035273515 | 0.3 | 0 | 0.259259 | T | C | AFUN005498 | UGT314A3 | c.17T>C | p.Leu6Pro | FUMOZ & FANG |
| 37737524 | 1.82E-07 | 1.10E-05 | 0.7 | 0 | 0.68254 | C | T | AFUN005498 | UGT314A3 | c.848C>T | p.Thr283Met | FUMOZ & FANG |
| 37739021 | 6.34E-12 | 1.59E-09 | 0 | 0.95 | 0.947368 | T | A | AFUN005498 | UGT314A3 | c.1117T>A | p.Phe373Ile | FUMOZ & FANG |
| 82048691 | 0.00137109 | 0.022661345 | 0.35 | 0 | 0.301587 | G | A | AFUN016158 | UGT301A3 | c.28G>A | p.Val10Ile | FUMOZ & FANG |
| 82049426 | 0.00234051 | 0.035273515 | 0.7 | 1 | 0.259259 | A | G | AFUN016158 | UGT301A3 | c.637A>G | p.Ile213Val | FUMOZ & FANG |
| 82049618 | 0.00234081 | 0.035273515 | 0.3 | 0 | 0.259259 | T | C | AFUN016158 | UGT301A3 | c.752T>C | p.Val251Ala | FUMOZ & FANG |
| 82050166 | 0.00233702 | 0.035273515 | 0.3 | 0 | 0.259259 | A | T | AFUN016158 | UGT301A3 | c.1300A>T | p.Thr434Ser | FUMOZ & FANG |
| 82056015 | 9.22E-10 | 1.05E-07 | 0 | 0.85 | 0.836601 | G | A | AFUN004354 | UGT301C2 | c.16G>A | p.Val6Ile | FUMOZ & FANG |
| 82056127 | 3.49E-05 | 0.000997223 | 0 | 0.5 | 0.455556 | A | T | AFUN004354 | UGT301C2 | c.128A>T | p.His43Leu | FUMOZ & FANG |
| 82056845 | 7.80E-07 | 3.80E-05 | 0.65 | 0 | 0.632479 | A | G | AFUN004354 | UGT301C2 | c.772A>G | p.Ile258Val | FUMOZ & FANG |
| 82057629 | 8.20E-07 | 3.92E-05 | 0.65 | 0 | 0.632479 | A | G | AFUN004354 | UGT301C2 | c.1556A>G | p.Asn519Ser | FUMOZ & FANG |
| 82057635 | 2.68E-06 | 0.000119946 | 0.6 | 0 | 0.574074 | G | A | AFUN004354 | UGT301C2 | c.1562G>A | p.Arg521Lys | FUMOZ & FANG |
| 82057646 | 3.03E-06 | 0.000122941 | 0.6 | 0 | 0.574074 | C | A | AFUN004354 | UGT301C2 | c.1573C>A | p.Pro525Thr | FUMOZ & FANG |
| 87996136 | 0.00187149 | 0.0302876 | 0 | 0.3 | 0.277778 | A | C | AFUN019724 | UGT308G4 | c.949A>C | p.Asn317His | FUMOZ & FANG |
| 87996328 | 0.000874523 | 0.015288292 | 0 | 0.35 | 0.31746 | C | T | AFUN019724 | UGT308G4 | c.1141C>T | p.His381Tyr | FUMOZ & FANG |
| 87996485 | 0.00234127 | 0.035273515 | 0 | 0.3 | 0.277778 | A | G | AFUN019724 | UGT308G4 | c.1298A>G | p.Tyr433Cys | FUMOZ & FANG |
| 87999732 | 0.000874523 | 0.015288292 | 0 | 0.35 | 0.31746 | G | A | AFUN009064 | UGT308G3 | c.1276G>A | p.Gly426Ser | FUMOZ & FANG |
| 88002475 | 0.00151661 | 0.024925642 | 0.7 | 1 | 0.259259 | A | G | AFUN004976 | UGT308G2 | c.1214A>G | p.Lys405Arg | FUMOZ & FANG |
| 90805096 | 0.00331652 | 0.047657657 | 0.35 | 0.8 | 0.303202 | A | T | AFUN003593 | UGT36C3 | c.1242T>A | p.Asp414Glu | FUMOZ & FANG |
| 381904 | 0.0153301 | 0.999879513 | 0 | 0.2 | 0.166667 | C | T | AFUN005786 | UGT306C2 | c.1367G>A | p.Arg456Lys | Malawi Dead & Alive |
| 382326 | 0.0382058 | 0.999879513 | 0.15 | 0 | 0.111111 | A | T | AFUN005786 | UGT306C2 | c.945T>A | p.His315Gln | Malawi Dead & Alive |
| 383945 | 0.0382698 | 0.999879513 | 0.15 | 0 | 0.111111 | C | A | AFUN016302 | UGT306A3 | c.1539G>T | p.Gln513His | Malawi Dead & Alive |
| 3258399 | 0.0382058 | 0.999879513 | 0 | 0.15 | 0.111111 | T | C | AFUN002058 | UGT49A4 | c.284T>C | p.Ile95Thr | Malawi Dead & Alive |
| 3258874 | 0.0382058 | 0.999879513 | 0.15 | 0 | 0.111111 | C | T | AFUN002058 | UGT49A4 | c.685C>T | p.Leu229Phe | Malawi Dead & Alive |
| 3259038 | 0.0382058 | 0.999879513 | 0.15 | 0 | 0.111111 | A | T | AFUN002058 | UGT49A4 | c.849A>T | p.Gln283His | Malawi Dead & Alive |
| 24375446 | 0.0382058 | 0.999879513 | 0 | 0.15 | 0.111111 | T | C | AFUN019845 | UGT302A3 | c.1507A>G | p.Ile503Val | Malawi Dead & Alive |
| 24376445 | 0.0204054 | 0.999879513 | 0.8 | 0.45 | 0.192872 | G | A | AFUN019845 | UGT302A3 | c.650C>T | p.Thr217Ile | Malawi Dead & Alive |
| 24376761 | 8.01E-05 | 0.999879513 | 0.6 | 0.05 | 0.500942 | G | C | AFUN019845 | UGT302A3 | c.334C>G | p.Gln112Glu | Malawi Dead & Alive |
| 24377150 | 0.0204054 | 0.999879513 | 0.8 | 0.45 | 0.192872 | C | T | AFUN019845 | UGT302A3 | c.83G>A | p.Gly28Asp | Malawi Dead & Alive |
| 37735579 | 0.024579 | 0.999879513 | 0.1 | 0.4 | 0.193122 | A | G | AFUN005498 | UGT314A3 | c.422A>G | p.His141Arg | Malawi Dead & Alive |
| 82056028 | 0.0382058 | 0.999879513 | 0 | 0.15 | 0.0740741 | G | T | AFUN004354 | UGT301C2 | c.29G>T | p.Cys10Phe | Malawi Dead & Alive |
| 82056845 | 0.0153301 | 0.999879513 | 0.2 | 0 | 0.166667 | A | G | AFUN004354 | UGT301C2 | c.772A>G | p.Ile258Val | Malawi Dead & Alive |
| 82057646 | 0.0442079 | 0.999879513 | 0.5 | 0.2 | 0.155556 | C | A | AFUN004354 | UGT301C2 | c.1573C>A | p.Pro525Thr | Malawi Dead & Alive |
| 3258498 | 0.0382058 | 0.806374302 | 0.15 | 0 | 0.111111 | A | G | AFUN002058 | UGT49A4 | c.383A>G | p.Gln128Arg | Uganda Dead & Alive |
| 24376011 | 0.0382058 | 0.806374302 | 0 | 0.15 | 0.111111 | T | A | AFUN019845 | UGT302A3 | c.1024A>T | p.Thr342Ser | Uganda Dead & Alive |
| 24376680 | 0.0382058 | 0.806374302 | 0 | 0.15 | 0.111111 | T | A | AFUN019845 | UGT302A3 | c.415A>T | p.Thr139Ser | Uganda Dead & Alive |
| 82056015 | 0.00234051 | 0.806374302 | 0 | 0.3 | 0.259259 | G | A | AFUN004354 | UGT301C2 | c.16G>A | p.Val6Ile | Uganda Dead & Alive |
| 90810470 | 0.0382058 | 0.806374302 | 1 | 0.85 | 0.111111 | T | C | AFUN003593 | UGT36C3 | c.226A>G | p.Thr76Ala | Uganda Dead & Alive |

**Table S9.** Primers used to validate the expression profiles of select UGTs using quantitative real-time PCR.

| **Vectorbase ID** | **Gene Name** | **Forward Sequence** | **Reverse Sequence** |
| --- | --- | --- | --- |
| AFUN006766 | RpS17 | AGAGGTGGCCATCATTCCGA | TCCTCCTGGAGCTTGATCGA |
| AFUN004354 | UGT301C2 | TTTGGCGAGAAGCTCCAAAAC | ACAGCTTGACGAAATCCGAT |
| AFUN011266 | UGT310B2 | TACTGCCACCGAACGTGATG | CGAAGCTAAACAGCACCACAC |

References

1.	Wondji, C.S., et al., *RNAseq-based gene expression profiling of the Anopheles funestus pyrethroid-resistant strain FUMOZ highlights the predominant role of the duplicated CYP6P9a/b cytochrome P450s.* G3 Genes|Genomes|Genetics, 2021. **12**(1).

2.	Wondji, C.S., et al., *Two duplicated P450 genes are associated with pyrethroid resistance in Anopheles funestus, a major malaria vector.* Genome Res, 2009. **19**(3): p. 452-9.

3.	Mugenzi, L.M.J., et al., *Cis-regulatory CYP6P9b P450 variants associated with loss of insecticide-treated bed net efficacy against Anopheles funestus.* Nat Commun, 2019. **10**(1): p. 4652.

4.	Mugenzi, L.M.J., et al., *A 6.5-kb intergenic structural variation enhances P450-mediated resistance to pyrethroids in malaria vectors lowering bed net efficacy.* Molecular Ecology, 2020. **29**(22): p. 4395-4411.

5.	Weedall, G.D., et al., *A cytochrome P450 allele confers pyrethroid resistance on a major African malaria vector, reducing insecticide-treated bednet efficacy.* Sci Transl Med, 2019. **11**(484).

6.	Ibrahim, S.S., et al., *The P450 CYP6Z1 confers carbamate/pyrethroid cross-resistance in a major African malaria vector beside a novel carbamate-insensitive N485I acetylcholinesterase-1 mutation.* Mol Ecol, 2016. **25**(14): p. 3436-52.

7.	Hearn, J., et al., *Multi-omics analysis identifies a CYP9K1 haplotype conferring pyrethroid resistance in the malaria vector Anopheles funestus in East Africa.* Mol Ecol, 2022. **31**(13): p. 3642-3657.

8.	Riveron, J.M., et al., *Genome-Wide Transcription and Functional Analyses Reveal Heterogeneous Molecular Mechanisms Driving Pyrethroids Resistance in the Major Malaria Vector Anopheles funestus Across Africa.* G3 Genes|Genomes|Genetics, 2017. **7**(6): p. 1819-1832.

9.	Riveron, J.M., et al., *A single mutation in the GSTe2 gene allows tracking of metabolically based insecticide resistance in a major malaria vector.* Genome biology, 2014. **15**(2): p. R27-R27.
